# Highly Accurate Cancer Phenotype Prediction with AKLIMATE, a Stacked Kernel Learner Integrating Multimodal Genomic Data and Pathway Knowledge

**DOI:** 10.1101/2020.07.15.205575

**Authors:** Vladislav Uzunangelov, Christopher K. Wong, Joshua M. Stuart

## Abstract

Advancements in sequencing have led to the proliferation of multi-omic profiles of human cells under different conditions and perturbations. In addition, several databases have amassed information about pathways and gene “signatures” – patterns of gene expression associated with specific cellular and phenotypic contexts. An important current challenge in systems biology is to leverage such knowledge about gene coordination to maximize the predictive power and generalization of models applied to high-throughput datasets. However, few such integrative approaches exist that also provide interpretable results quantifying the importance of individual genes and pathways to model accuracy. We introduce AKLIMATE, a first kernel-based stacked learner that seamlessly incorporates multi-omics feature data with prior information in the form of pathways for either regression or classification tasks. AKLIMATE uses a novel multiple-kernel learning framework where individual kernels capture the prediction propensities recorded in random forests, each built from a specific pathway gene set that integrates all omics data for its member genes. AKLIMATE outperforms state-of-the-art methods on diverse phenotype learning tasks, including predicting microsatellite instability in endometrial and colorectal cancer, survival in breast cancer, and cell line response to gene knockdowns. We show how AKLIMATE is able to connect feature data across data platforms through their common pathways to identify examples of several known and novel contributors of cancer and synthetic lethality.

## Introduction

The drop in sequencing cost has made it common for biological experiments to generate multiomic profiles under a variety of conditions and perturbations. For example, the Cancer Genome Atlas (TCGA) contains thousands of patient samples with simultaneous copy number, mutation, methylation, mRNA, miRNA and protein levels measurements [33]. The analysis of multi-omic experiments produces feature sets that capture the genomic and transcriptomic changes relevant to a specific condition, sample subtype, or biological pathway - this “prior knowledge” eventually accumulates in a growing number of databases [12][47][18][88][58]. However, the integration of such prior knowledge in the analysis of multi-omic data from new experiments remains a significant challenge that is still not fully solved.

There are three main challenges that are inherent to bioinformatics algorithms for supervised and unsupervised learning - prior knowledge integration, heterogeneous data interrogation, and ease of interpretation. New methods try to address some aspects of these three main challenges (for recent reviews, see [60][49][35]). Several common approaches emerge for supervised learning of sample phenotypes. A popular way of increasing interpretability is to train models with regularization terms that constrain the number of included features - sparse models are considered easier to interpret than ones containing thousands of variables. Common regularization penalties are the lasso [76] and the elastic net [87]. More sophisticated regularization schemes can control the model behavior at the feature set level (e.g. the group lasso (GL) [86] and the overlap group lasso (OGL) [37]), allowing the incorporation of prior knowledge in the form of feature sets. However, both have drawbacks that make them less suitable for biological analysis where genes often participate in multiple processes. GL requires that if a feature’s weight is zero in one group, its coefficients in all other groups must necessarily be zero. OGL tends to assign positive coefficients to entire feature sets, making it less suitable in situations where the latter are noisy or only partially relevant.

Network-regularized methods [69][46] represent a different approach where gene-gene interaction networks are used as regularization terms on the L_2_-norm of feature weights. While they do use pathway-level information and are straightforward to interpret, they generally focus on an individual data type.

Multiple Kernel Learning (MKL) approaches [3][59][74][27][66] [50] can incorporate heterogeneous data by mapping each set of features through a kernel function and learning a linear combination of the kernel representations. Each kernel represents distinct sample-sample similarities providing flexible and powerful transformations to access either explicit or implicit feature combinations. To prevent overfitting, MKL methods generally include a regularization term - e.g., L_1_ sparsity-inducing norm on the kernel weights [59] or the elastic net [74]. Prior knowledge can also be integrated by constructing individual kernels from a pathway’s member features within each data type [29][27][66][78][50]. Indeed, MKL methods with prior knowledge integration [29][14] have won several Dialogue on Reverse-Engineering Assessment and Methods (DREAM)[70] challenges, including a predecessor of the approach described here [78][29]. Nevertheless, MKL suffers significant drawbacks when the contributions of input features need to be evaluated. Except in trivial cases, it is generally impossible to assign importance to the original features once the method is trained in the kernel function feature space, thus limiting interpretability of solutions. In addition, feature heterogeneity necessitates the construction of separate kernels for each data type, limiting the ability of MKL to capture cross-data-type interactions.

All these methods (and more) can be used as components in more complex ensemble learning models. Ensembles combine predictions from multiple algorithms into a more robust, and often more accurate, “wisdom of crowds” final prediction. The simplest and most common ensemble technique is averaging the predictions of component models. Averaging over uncorrelated models or models with complementary information can improve performance - it is one of the main reasons for the emergence of collaborative competitions such as the DREAM challenges [51][14]. In fact, such ensemble methods often win DREAM challenges [13] or out-perform competitors in genomic prediction tasks [41]. While they provide a boost in predictive accuracy, their interpretation is quite challenging - ensembles often combine different model types, making the computation of input feature importance impossible in the large majority of cases. A notable exception is the Random Forest (RF)[8] - an ensemble of decision trees that has been widely applied to bioinformatics problems.

A more general approach to combining the predictions of multiple models is model stacking [84]. In stacking, component (base) learner predictions are used as inputs to an overall (stacked) model which produces the final calls. To reduce overfitting and improve generalization, base learner predictions are generated in a cross-validation manner - a sample’s predicted label comes from a model trained on all folds except the one that includes the sample in question. Importantly, if the stacked model is a weighted combination of the base models (a Super Learner), it is asymptotically guaranteed to perform at least as well as the best base learner or any conical combination of the base learners [81][57]. Stacked models exhibit identical pros and cons as standard ensemble models, with diminished interpretability traded off for increased accuracy. In some cases, however, interpretability can be tractable - e.g. [82] uses RF base learners with a stacked least square regression to compute a final weighted average of the base RF predictions. Although not examined in [82], we demonstrate that a similar setup can lead to an intuitive derivation of feature importance scores.

In this study we introduce the Algorithm for Kernel Learning with Integrative Modules of Approximating Tree Ensembles (AKLIMATE) – a novel approach that combines heterogeneous data with prior knowledge in the form of gene sets. AKLIMATE can evaluate the predictiveness of individual features (e.g. genes) as well as feature sets (e.g. pathways). It harnesses the advantages of RFs (native handling of continuous, categorical and count data, invariance to monotonic feature transformations, ease of feature importance computation), MKL (intuitive integration of overlapping feature sets), and stacked learning (improved accuracy) while avoiding many of their shortcomings. AKLIMATE relies on three major computational insights. First, we use summary statistics of decision trees within an RF model to compute a kernel similarity matrix (RF kernel) that is data driven yet capable of capturing complex non-linear feature relationships. Second, if an RF model is trained with only the features mapping to a distinct biological process, signature or genomic location, converting that model to an RF kernel allows us to handle many different kinds of heavily overlapped feature groups without the undesirable side effects of GL/OGL. Third, the RF kernel naturally re-weights the input features so that more informative ones make bigger contributions to kernel construction. This final property is key when dealing with noisy or partially relevant feature set definitions.

We first provide a detailed explanation of the AKLIMATE methodology. Then, we demonstrate that AKLIMATE outperforms state-of-the-art algorithms on various classification and regression tasks - microsatellite instability in endometrial and colon cancer, survival in breast cancer, and small hairpin RNA (shRNA) knockodown viability in cancer cell lines.

## Methods

### Overview

Mock figure for multimodal data adapted from [83]. Mock figure for pathway databases adapted from Pathway Commons website. Mock figure for methylation of a genomic region adapted from Wikipedia.

AKLIMATE is a stacked learning algorithm with RF base learners and an MKL meta-learner. Each base learner is trained using only the features in a particular feature set that corresponds to some known biological concept or process - e.g. a biological pathway, a chromosomal region, a drug response or shRNA knockdown signature, or disease subtype biomarkers (Fig.1). Each feature in a feature set can be associated with multiple data modalities – i.e. a gene will have individual features corresponding to its copy number, mutation status, mRNA expression level, or protein abundance. Furthermore, although feature sets normally consist of genes, they are easily extendable to a more comprehensive membership. For example, they can be augmented with features for mutation hotspots, different splice forms, or relevant miRNA or methylation measurements. AKLIMATE’s MKL meta-learner trains on RF kernel matrices (see the *RF Kernel* section), each of which is derived from a corresponding base learner (Fig.2). An RF kernel captures a proximity measure between training samples based on the similarity of their predicted labels and decision paths in the trees of the RF. The MKL learning step finds the optimal weighted combination of the RF kernels. The MKL optimal solution can be interpreted as the meta-kernel associated with the most predictive meta-feature set derived from all interrogated feature sets.

**Fig. 1.**
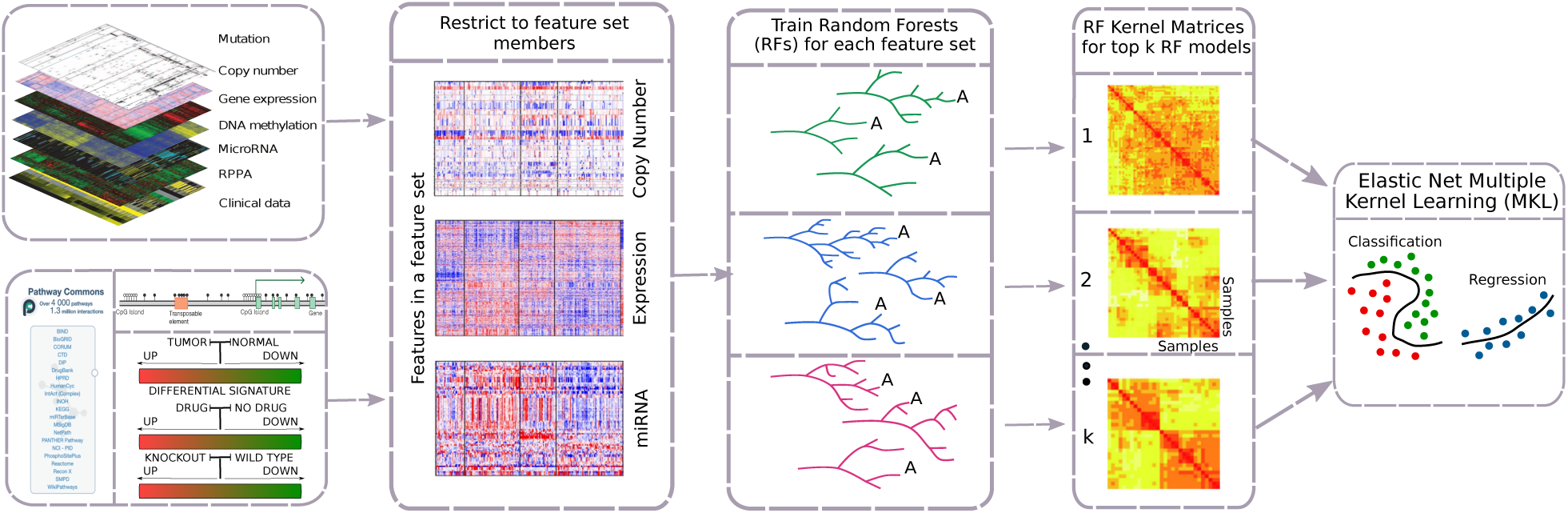
Overview of AKLIMATE. AKLIMATE takes as inputs multiple data types and a collection of feature sets. AKLIMATE first trains RF base learners, one for each feature set, with all available multi-omic features that map to the set in question. The RFs are then ranked by their predictive performance and the top K are converted to RF kernels. Finally, the RF kernels are used as input in an elastic net MKL meta-learner to produce the final predictions. Elastic net hyperparameters are optimized via cross-validation.

**Fig. 2.**
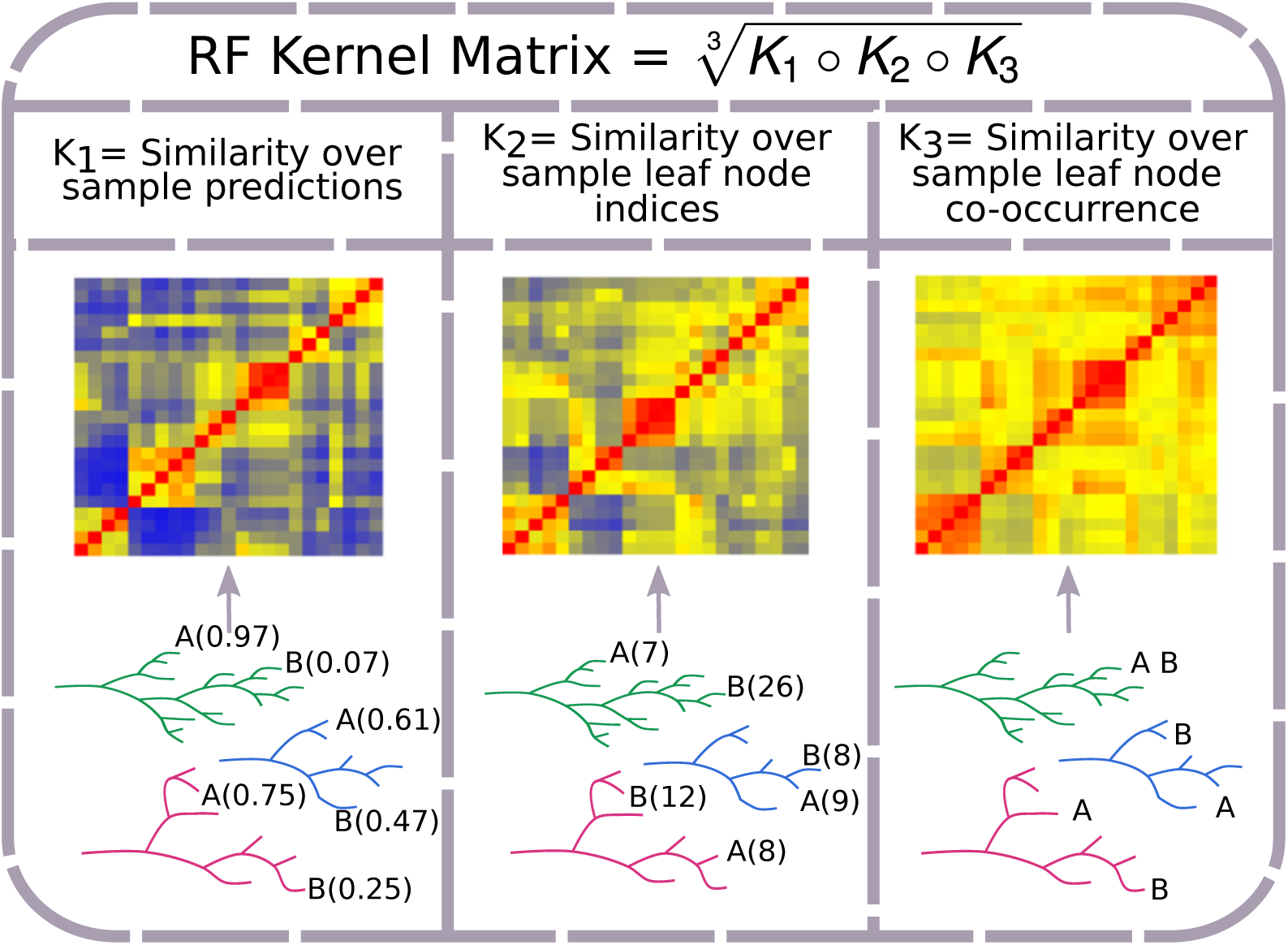
RF kernel matrix construction. The RF Kernel matrix is a geometric mean of the Hadamard product of three component similarity matrices. Each component captures a different aspect of a RF model. *K*_1_ captures the similarity over RF tree predictions for sample labels (in the case of classification, probability of belonging to a class); two samples (A and B) show different predicted probabilities of belonging to the positive class (probability estimate in parentheses). *K*_2_ represents the similarity over RF tree leaf indices to which samples are assigned; predicted leaf indices shown in parentheses for the two samples. *K*_3_ reflects the proportion of times samples are assigned to the same RF tree leaf; e.g. the two samples end up together in one out of three example trees.

The key contributions of AKLIMATE are

1. The introduction of an integrative empirical kernel function that combines similarity in predicted labels with proximity in the space of RF trees (RF kernel).
2. The extension of kernel learning to a stacking framework. To our knowledge, AKLIMATE is the first stacked learning formulation to incorporate base kernels.

In the next sections we give a detailed description of each AKLIMATE component.

### Random Forest

An RF learner is an ensemble of decision trees each of which operates on a perturbed version of the training set. The perturbation is achieved by two randomization techniques - on one hand, each tree trains on a bootstrapped data set generated by drawing samples with replacement. On the other hand, at each node in a decision tree, only a subset of all features are considered as candidates for the next split. Tree randomization tends to reduce correlation among predictions of individual trees at the expense of increased variance of error estimates [48]. However, due to the averaging effect of ensembles of decorrelated models, it generally reduces the overall error variance of the ensemble RF [48]. That effect is partially negated by an increase in prediction bias, but broadly speaking the more decorrelated the trees the better the RF performance [48].

As each tree is constructed from a subsample of the training data, there is a tree-specific set of excluded, or out-of-bag (OOB) samples. The OOB predictions for the training set of a regression task are computed by walking the OOB samples along their respective trees and averaging the leaf OOB predictions for each sample across the forest:

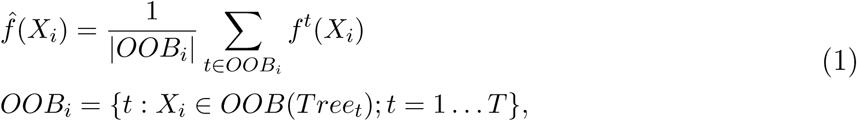

where *X*_*i*_ are the features associated with sample *i, OOB*_*i*_ is the set of trees in which sample *i* is OOB, *T* is the total number of trees in an RF and *f*^*t*^(*X*_*i*_) is the classification of the *i*^*th*^ sample in tree *t*. Similarly, for classification tasks the averaging of tree OOB predictions is replaced by majority vote.

It has been shown that the OOB error rate provides a very good approximation to generalization error [8]. For that reason, AKLIMATE uses RF kernels based on OOB predictions as level-one data for the MKL meta-learner training (see *RF Kernel* and *Stacked Learning* below).

### Kernel Learning

Kernel learning is an approach that allows linear discriminant methods to be applied to problems with non-linear decision boundaries [64]. It relies on the “kernel trick” - a transformation of the input space to a different (potentially infinite dimensional) feature space where the train set samples are linearly separable. The utility of the “kernel trick” stems from the fact that the feature map between the input space and the feature space does not need to be explicitly stated - if the kernel function is positive definite (i.e. it has an associated Reproducing Kernel Hilbert Space), we only need to know the functional form of the feature space dot product in terms of the input space variables [64][2]. A kernel function represents such a generalized dot product.

Common approaches construct kernel functions using an explicit closed form, such as a polynomial or a radial basis function. Kernel functions that encode more complex relationships between training objects also exist, particularly if the objects can be defined in a recursive manner (ANOVA kernels, string kernels, graph kernels, see [68]). However, knowing the closed form kernel function is not a necessary condition - if the kernel positive definiteness can be verified and the kernel matrix of pairwise dot products (similarities) can be computed, we can still utilize the “kernel trick”. In fact, the Representer Theorem [42][64] guarantees that the optimal solution to a large collection of optimization problems can be computed by kernel matrix evaluations on the finite-dimensional training data, ensuring the applicability of kernel-based alogirthms to real-world problems. More formally, for an arbitrary loss function *L*(*f* (*x*_1_) … *f* (*x*_*N*_)), the minimizer *f* * of the regularized risk function

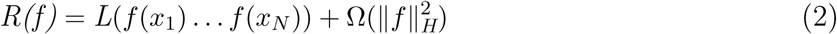

can be expressed as

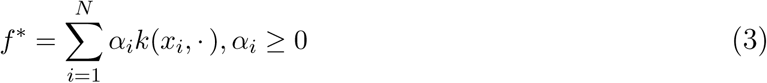

provided that *k*(*·, x*) is a positive definite kernel and the regularization term 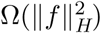 is a monotonically increasing function.

AKLIMATE uses the dependency structure of a RF trained on a, possibly multi-modal, feature set (e.g. a pathway) to define an implicit empirical kernel function. It then computes an RF kernel matrix (see *RF Kernel* below) of pairwise training set similarities to use in a kernel learning algorithm. As different feature sets can contribute complementary information for a given prediction task, AKLIMATE utilizes a Multiple Kernel Learning approach that can integrate the associated RF kernels into an optimal predictor.

### Multiple Kernel Learning

Kernel learning can lead to large improvements in accuracy, provided that an optimal kernel is selected. That choice can be difficult, however - the best kernel function for a heterogeneous training data set may be non-obvious, may entail tuning several hyperparameters, or may even lack a closed form expression. MKL addresses the problem of kernel selection by constructing a composite kernel that is a data-driven optimal combination of candidate kernels [3][44][28][43]. Linear combinations of kernels of the form 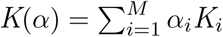 with either a conical (*∀i, α* _*i*_ ≥ 0) or convex 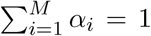 sum constraint on the kernel weights are by far the most commonly used because of their many desirable properties (although algorithms that allow non-linear combinations do exist - see [28]). Some important advantages are:

1. Conical/convex combinations of positive definite kernels are positive definite.
2. The composite kernel is associated with a feature space that is the concatenation of all individual kernel feature spaces.
3. The Representer Theorem is readily extendable to the the conical/convex combination case - the optimal solution *f* * takes the form 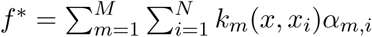 [74][64].

MKL algorithms admit different forms of regularization depending on what norm of the kernel weights is chosen. AKLIMATE uses an elastic-net [87] regularizer on the norm of the individual kernel weights of the form 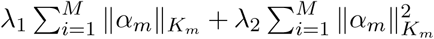, where *λ*_1_, *λ*_2_ *≥* 0, *α*_*m*_ = (*α*_*m*,1_ … *α*_*m,N*_)^*T*^, and 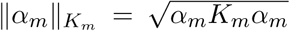 is the definition of a kernel norm as in [74] and [73]. The elastic net regularization provides the flexibility to find both sparse and dense solutions with appropriately tuned *λ*’s – i.e. *λ*_1_ *>> λ*_2_ gives few non-zero kernel weights (sparse) while *λ*_1_ *< λ*_2_ gives many non-zero kernel weights (dense). Also note that *λ*_1_ = 0 and *λ*_2_ *>* 0 allows for a uniform kernel weight solution.

The explicit form of AKLIMATE’s optimization problem is:

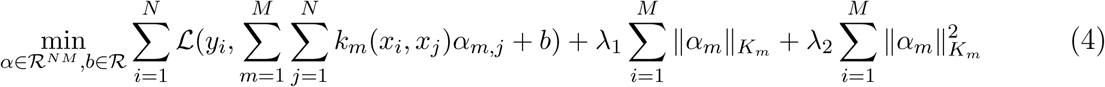

and is solved using the SpicyMKL algorithm described in [74]. The optimal kernel weights are recovered by:

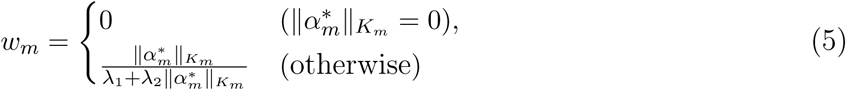

where *α*\* is the solution to (4). The model weights are re-scaled to satisfy 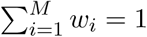.

### RF Kernel

The most common approach to kernel matrix evaluation is to specify an explicit data dependency model for the kernel function - e.g. polynomial, Gaussian, ANOVA or graph kernels [68]. However, choosing the dependency structure a priori can lead to lack of robustness, particularly when it is not obvious what the right dependency is. AKLIMATE implements a data-driven framework that approximates the true dependency model by means of a RF and evaluates its implicit kernel function (the RF kernel). RF kernels are robust to overfitting, generalize well to new data, and facilitate the integration of signals across data types.

Defining a kernel through a RF is not a new idea - in fact the concept was introduced at the same time as RFs [7]. In particular, for a random forest of trees with equal number of leaves and uniform predictions in each leaf (e.g. fully grown trees with leaf size of 1), [7] defines a positive definite kernel based on the probability two samples share a leaf:

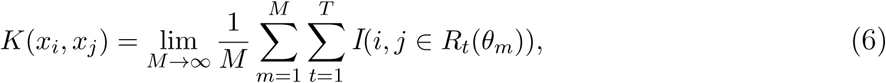

where *M* is the number of trees in the forest, *T* is the number of leaves in a tree, *θ*_*m*_ is a variable capturing the random selection of training set samples and features in the process of constructing the *m*th tree, *R*_*t*_ is the *t*th leaf of the *m*th tree, and *I*(·) is the indicator function. The finite approximation of Eqn.6 (*M < ∞*; Fig.2, *K*_3_ kernel) is positive semi-definite [20]. Kernels generated from RFs and their theoretical properties are also discussed in [25] and [65].

AKLIMATE’s RF kernel extends the original definition (Eqn. 6) by incorporating two additional RF-derived statistics. The first intuition is that two samples predicted to have the same label across the trees of an RF should be considered alike even if they happen to fall into different leaves (Fig.2, *K*_1_ kernel). The second intuition is that earlier node splits in a tree generally separate more distinct sample groups, while later splits tend to fine tune the decision boundary, highlighting more subtle differences. Thus, irrespective of the predicted labels, two samples that end up in different tree depths – one early- and one late-split leaf – should be considered less similar than samples landing in two late-split leaves (Fig.2, *K*_2_ kernel). Incorporating these three patterns, AKLIMATE’s RF kernel matrix is computed using the following steps:

1. Calculate similarity over predictions across RF trees using:

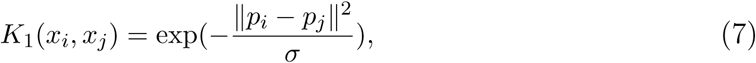

where *p*_*i*_ = (*p*_*i*,1_, *p*_*i*,2_, …, *p*_*i,M*_) is a vector of predictions for data point *i* from the *M* trees in the RF. *p*_*i*_ always has continuous entries - either the actual predictions in a regression setting, or the probabilities of class membership for classification problems. The | |*p*_*i*_ *− p*_*j*_| |^2^ represent distances between prediction vectors and are divided by the scaling constant *σ* = max_*i,j*_ | |*p*_*i*_ *− p*_*j*_| |^2^ so that they lie in the [0, 1] range. Exponentiation of the negative distances converts them to similarities.
2. Compute similarity over leaf node indices across RF trees:

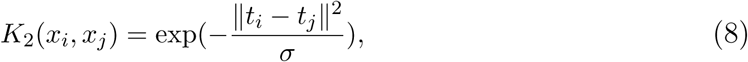

where *t*_*i*_ = (*t*_*i*,1_, *t*_*i*,2_, …, *t*_*i,M*_) is a vector of the leaf indices of sample *i* across the RF. Tree nodes are indexed starting from the base node and then sequentially across each depth level of the tree. The scaling constant *σ* is set in the same manner as in *K*_1_.
3. Calculate similarity as the frequency of leaf node co-occurrence using the finite approximation of Eqn.(6) with variable number of leaves per tree:

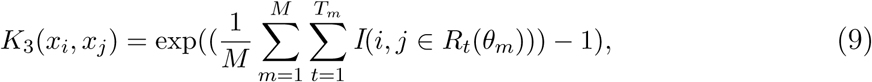

where the exponential transformation is added to keep *K*_3_ on a similar scale to *K*_1_ and *K*_2_ in order to avoid any one component having a disproportionate effect.
4. Calculate the RF kernel as the geometric mean of the element-wise product of *K*_1_,*K*_2_ and *K*_3_:

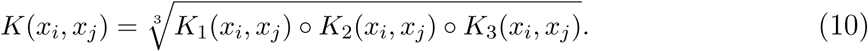

Importantly, since the components *K*_1_, *K*_2_, and *K*_3_ are positive definite, so too is *K. K*_1_ and *K*_2_ are Gaussian kernels over *𝒫 × 𝒫, 𝒫* ⊆ *ℑ*^+^ and *𝒯 × 𝒯, 𝒯* ⊆ *ℑ*^+^ respectively, which ensures their positive definiteness [68]. *K*_3_ is the exponent of a positive-definite kernel, i.e. a positive-definite kernel as well [68]. Finally, *K* is the Hadamard product of three positive-definite kernels raised to a positive power – both of these operations preserve positive-definiteness [68].

### Stacked Learning

Stacked learning is a generalization of ensemble learning in which the stacked model (meta-learner) uses the prediction output of its components (base learners) as training data for the computation of the final predictions [84][6]. What makes stacked learning unique is that the base learner predictions (level-one data) are generated in a cross-validated manner that excludes each sample from the training set that produces its predicted label. More specifically, if our training set (level-zero data) is *X* = *{X*_*i*_ : *i* = 1 … *N, X*_*i*_ *∈ ℛ*^*p*^} with labels *Y* = *{Y*_*i*_ : *i* = 1, …, *N, Y*_*i*_ *∈ ℛ}* and we have a collection of base learners **Δ** = {Δ_1_, …, Δ_*S*_} then the stacked generalization proceeds as follows [57][45]:

1. Randomly split the level-zero data into *V* folds of roughly equal size - *h*_1_, …, *h*_*V*_ (*V* -fold cross validation). Note that each *h*_*v*_ defines a set of indices that select a subset of the samples in *X*.
2. For each Δ_*s*_ base learner, train *V* models, collectively denoted as 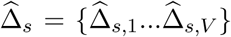 Each model 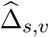 uses 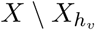 to train on and generates predictions for 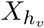.
3. Concatenate the *V* sets of predictions from 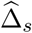 into a vector *z* of length *N*. The *N × S* matrix *Z* of such vectors for all *S* base learners becomes the feature matrix for level-one training.
4. Train a meta-learner Ψ on *Z* with hyperparameter tuning if necessary, again using *Y* as the labels.
5. The base learners used in the final stacked model are created using all training samples, so one final (non-cross-validated) training round is needed. To this end, train each Δ_*s*_ base learner on the full level-zero training set *X*.
6. Form the stacked model using the base-learners trained on the full data set and the meta-learner to obtain ({Δ_*s*_ : *s* = 1, …, *S*}, Ψ). To predict on a new data point *X*_*new*_, compute a 1 *× S* vector *Z*_*new*_ = {Δ_*s*_(*X*_*new*_) : *s* = 1, …, *S*} and use Ψ(*Z*_*new*_) as the stacked model’s prediction.

The predictive performance of the meta-learner is generally improved when the base learner predictions are maximally uncorrelated. This is achieved either by using different algorithms, or by varying the parameters of a particular modeling approach [6][81]. AKLIMATE generates diversity through the use of feature subsets built around distinct biological concepts and processes.

### Super Learner

Stacked learning imposes no conditions on the choice of meta-learner Ψ. The disadvantage of such flexibility is the lack of theoretical results for the improved empirical performance of stacking. A Super Learner [81] is a type of stacked learner with restrictions on Ψ that give provable desirable properties. The main such constraint is that the optimal meta-learner Ψ* is the minimizer of a bounded loss function. A Super Learner for which {Δ_1_, …, Δ_*S*_} and Ψ have uniformly bounded loss functions exhibits the “oracle” property - Ψ* is asymptotically guaranteed to perform as well as the optimal base learner 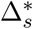 under the true data-generating distribution [80][79]. Furthermore, if we constrain the choice of Ψ to 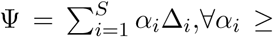, Ψ* asymptotically converges to the performance of the optimal conical combination of {Δ_1_, …, Δ_*S*_} [81][57]. This is the main reason why Ψ often takes the form of regularized linear or logistic regression.

For example, if the aim is to predict a continuous variable, one can set 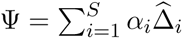 and solve the regression problem:

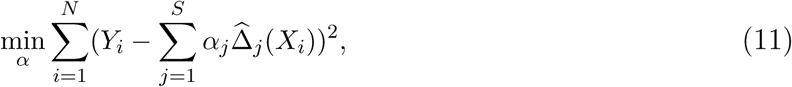

which can be regularized or subjected to a convex sum constraint on the *α* weights (e.g. 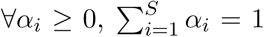 [81] [57]. Similarly, the squared error loss of (11) can be replaced with the logistic loss to solve a classification problem:

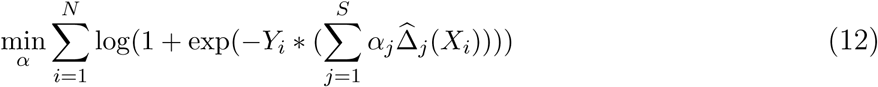

maintaining all theoretical Super Learner results.

### AKLIMATE

To our knowledge, AKLIMATE is the first instance of a kernel-based stacked learner. The base learners {Δ_1_, …, Δ_*S*_} are RFs, each of which is used to produce an associated kernel. The meta-learner Ψ is an elastic-net regularized MKL that can be interpreted as the kernel learning counterpart of linear regression. Before describing the algorithm, we first discuss how the level-one RF kernels are constructed using OOB samples.

### AKLIMATE Level-One (OOB) Kernel Construction

AKLIMATE uses RF kernels as the level-one training data. Normally, level-one data is generated with cross-validation to help the meta-learner avoid overfitting. Analogously, AKLIMATE utilizes out-of-bag (OOB) samples to generate the components of the RF kernels. For each pair of samples in the RF, the kernel similarity matrix is calculated using only those trees for which both of the samples have been withheld, with the following procedure:

1. For a RF with *M* trees, define *OOB*(*m*) as the set of samples that are OOB in the *m*th tree. Let *I*_*OOB*_ be the tree-level indexing function recording when both samples *i* and *j* are simultaneously OOB in a given RF; i.e.:

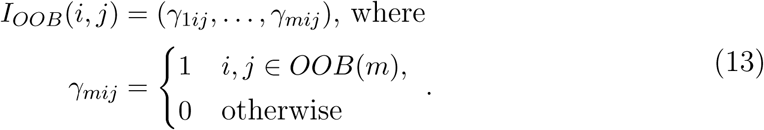
2. Compute the first constituent kernel matrix:

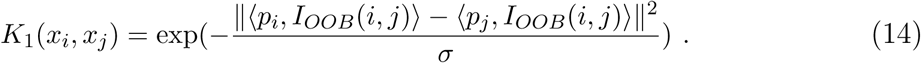
3. Compute the second constituent kernel matrix:

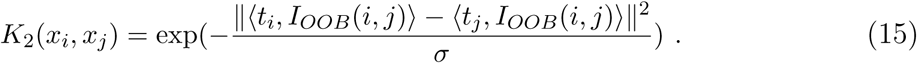
4. Compute the third constituent kernel matrix:

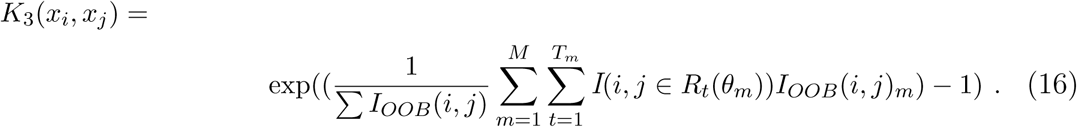
5. Calculate the combined kernel matrix:

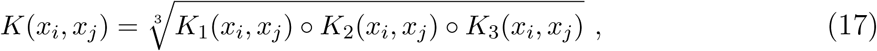

with the same notation and *σ* calculations as in (7)-(10).

AKLIMATE’s MKL meta-learner has two elastic-net hyperparameters (*λ*_1_, *λ*_2_) that require tuning. This is done by generating a random set of (*λ*_1_, *λ*_2_) pairs and ranking them based on *V* -fold (default *V* = 5) cross-validation fit with OOB RF kernels as input. To improve generalization, we use a simplified version of the overfit correction procedure in [56] - instead of selecting the hyperparameters that produce the best CV fit, we choose the ones corresponding to the 90th percentile of the distribution of the CV fit metric.

### AKLIMATE Algorithm

We next describe AKLIMATE’s learning algorithm. The method is given as input training data (*X, Y*) = {(*X*_*i*_, *Y*_*i*_) : *X*_*i*_ = *X*_*i*1_ ∪…∪ *X*_*id*_, *i* = 1 … *N, d* = 1 … *D*} consisting of *N* samples and *D* data types with feature memberships 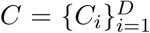 respectively. In addition, *S* feature sets (e.g. pathways) are supplied, each containing a list of features *P*_*s*_ such that the complete set is 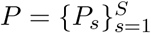. The algorithm outputs a meta-learner, Ψ* and a set of selected base learners **Δ**\*. In addition, a user-supplied parameter *G* determines the number of top feature sets (pathways) to incorporate for each sample during the base learning step. We use *G* = 5 in practice.

Formally, AKLIMATE operates according to the pseudocode in Algorithm 1. First AKLIMATE trains a separate RF for each feature set using data from all modalities relevant to the features in the set. Model accuracy *τ* is stored so that the top *L* models (by *τ*) can be selected that correctly predict an individual sample. A final set of relevant RFs **Δ**\* is obtained by taking the union over all sample-specific top *L* models. The helper function RFKERNELOOB creates a kernel from a given RF utilizing the out-of-bag approach described previously. MKL uses these level-one kernels to tune the elastic net hyperparameters *λ*_1_ and *λ*_2_ based on cross-validation performance. A final meta-learner Ψ* is then trained by running MKL with kernels constructed from the full RFs (RFKERNEL helper function) and with the determined elastic net hyperparameters.

#### Algorithm 1: AKLIMATE

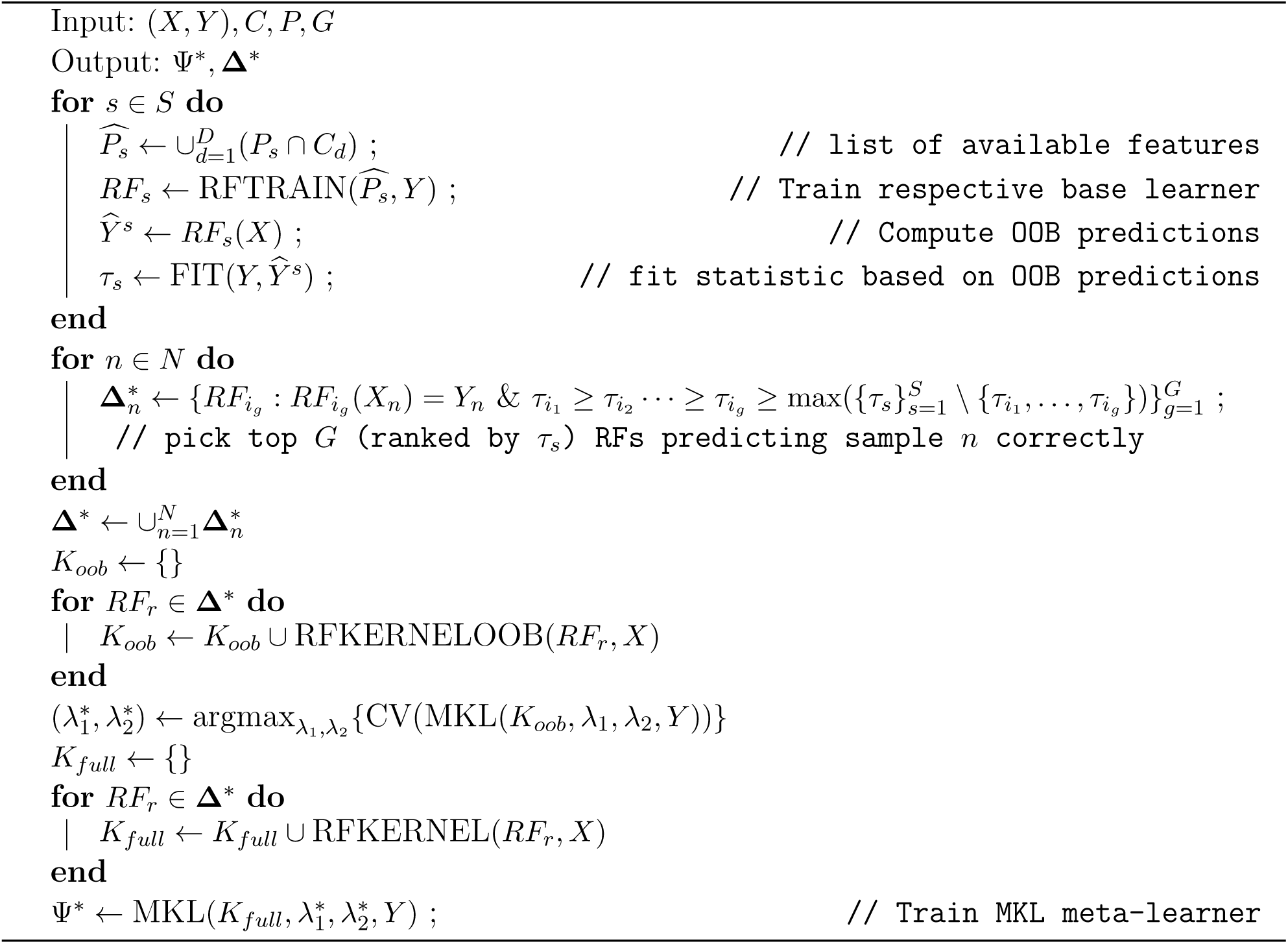

### AKLIMATE selection of relevant RFs

AKLIMATE’s selection step for the best RFs **Δ**\* in Algorithm 1 can filter the full collection of feature sets down to a subgroup of relevant sets two or more orders of magnitude smaller in size. This makes it possible to incorporate significantly more feature sets than standard MKL algorithms which require kernel evaluation for all feature sets. When the number of such sets is in the thousands, MKL can be computationally very slow, even for data sets of small sample size. However, usually only a small proportion of the feature set collection is truly explanatory for a given prediction task. Thus, filtering out the non-relevant parts of the compendium does not impact MKL accuracy yet drastically improves computation time.

Algorithm 1 demonstrates the **Δ**\* discovery process for a classification task. However, if *Y* is continuous (i.e. a regression problem), the predictions and labels are not directly comparable for equality. One approach is to take the squared error of the prediction-label differences and use that metric to re-rank RFs for each sample. In our experience, this leads to the selection of suboptimal RFs due to overfitting. Instead, AKLIMATE uses a more robust scheme that shows better results in practice - the vector of predictions for each sample across all 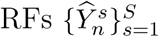 is binarized into matching and non-matching predictions and then the standard classification case selection rule is applied. The binarization is done as follows:

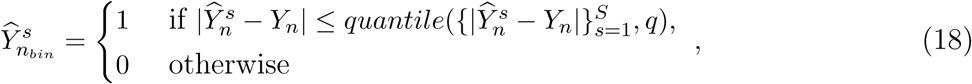

where *q* is a user-specified quantile of the empirical distribution of absolute prediction errors 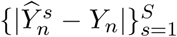 (default *q* = 0.05). This setup prioritizes RFs that perform near-optimally on individual data points and optimally when the training set is considered as a whole.

### AKLIMATE importance weighting of individual features and feature sets

Feature set weights *w*^*F S*^, 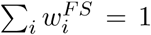, are recovered directly from the optimal MKL meta-learner Ψ*(Eqn.(5)). Feature weights can be calculated as 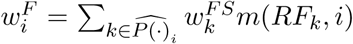 where 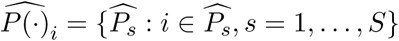 is the set of all feature sets that have feature *i* as its member, and *m*(*RF*_*k*_, *i*) is an *RF* -specific feature importance score computed from the *k*th *RF* with feature *i* among its input features.

The simplest way to compute *m*(*RF*_*k*_, *i*) is by averaging the improvement in the splitting criterion over all nodes that used feature *i* as a splitting variable. For the often used Gini impurity measure, this involves computing the mean difference in impurity before and after each split, with larger mean impurity decreases indicative of more important variables [9]. While fast, this rule suffers from important shortcomings - for example, it is biased in favor of variables with more potential split points (e.g. continuous or categorical with a large number of categories), particularly when trees train on bootstrapped data [71].

Many alternative *m*(*RF*_*k*_, *i*) rules have been proposed [71][62][1]. For our work, we choose as default the permutation-based importance calculation described in the original RF paper [8] - the vector of measurements for feature *i* is randomly permuted and the permuted variable is used in the calculation of OOB predictions; the difference in error rate between the permuted and non-permuted OOB predictions is taken as a measure of feature *i*’s importance. Permutation-based importance is robust and generally performs on par with more complex *m*(*RF*_*k*_, *i*) rules. Its biggest drawback is the higher computational cost. In cases where compute time is the main constraint, we recommend the actual impurity reduction (AIR) metric [55], which is an extension of the pseudodata-augmented approach in [62]. It is similar in speed to using the Gini impurity importance, but retains the desirable properties of permutation-based methods. While AIR can lead to a small prediction accuracy penalty, in our experience this effect has been negligible for classification tasks (for regression problems we still recommend permutation-based importance).

## Results

We evaluated AKLIMATE on multiple prediction tasks - microsatellite instability in endometrial and colon cancer, survival in breast cancer, and shRNA knockdown viability in cancer cell lines. We benchmarked AKLIMATE against comparable methods that have performed well in recent DREAM challenges. We chose both classification and regression tasks as well as various levels of data availability - a single data type, multiple data types (including inferred data), or multiple data types with clinical information.

### Microsatellite Instability

We first tested AKLIMATE on predicting microsatellite instability in the colorectal (COAD-READ) and endometrial (UCEC) TCGA cohorts. Microsatellite instability (MSI) arises as a result of defects in the mismatch repair machinery of the cell. Tumors with MSI (often accompanied by higher mutation rates) represent a clinically relevant disease subtype that is associated with better prognosis. MSI is also an immunotherapy indicator as such tumors produce more neoantigens. MSI can be predicted with high accuracy from expression alone [27], providing a straightforward benchmark for AKLIMATE performance with a single feature type in a binary classification setting.

We used expression data and MSI annotations for the COADREAD and UCEC TCGA cohorts. The UCEC cohort consisted of 326 patients, of which 105 exhibit high microsatellite instability (MSI-H) and the remaining 221 are classified as either low (MSI-L) or stable (MSS). The COADREAD cohort included 261 samples, with 37 MSI-H and the remaining 224 classified as either MSI-L or MSS. In both tumor types, we trained models to distinguish MSI-H patients from MSI-L+MSS patients on 50 phenotype-stratified partitions of 75% training and 25% test folds. We then computed area under the ROC curve (AUROC) for each set of test fold predictions.

We compared AKLIMATE to Bayesian Multiple Kernel Learning (BMKL) because it performed well in several DREAM challenges, in particular winning the NCI-DREAM Drug Sensitivity Prediction Challenge[14]. Furthermore, its pathway-informed extension [27] shares several similarities with AKLIMATE - it is a multiple kernel learning method that operates on pathway-derived kernels. In particular, BMKL uses expression-based Gaussian kernels computed on features from the PID pathway collection [63]. We tested four versions of BMKL - sparse single-task BMKL(SBMKL), dense single-task BMKL(DBMKL), sparse multi-task BMKL (SBMTMKL), and dense multi-task BMKL (DBMTMKL). Sparse BMKL models are the focus of [27] - they use sparsity-inducing priors to train models with few non-zero kernel weights (we used the hyperparameters specified in [27]). We added DBMKL and DBMTMKL to the comparison because dense MKL models (almost all kernels receive non-zero weights) tend to produce higher predictive accuracy in many experimental settings (e.g. [77]). Their parameters were identical to the ones for the sparse models except for (*ζ*_*κ*_, *η*_*κ*_), which were set to (999, 1) in the dense models ((*ζ*_*κ*_, *η*_*κ*_) = (1, 999) in the sparse ones). Finally, the single-task models were trained separately on the UCEC and COADREAD cohorts, while the multitask versions learned MSI status on the two TCGA cohorts jointly, with each cohort representing a separate task. All train/test splits were matched across methods. All BMKL models used 196 PID gene sets; model parameters, kernel computations and data filtering steps matched the setup in [27].

Since AKLIMATE uses a much larger gene set compendium (*S* = 17, 273 feature sets, see *Supplementary Information*), we controlled for this prior information imbalance as a possible source of performance bias. For that purpose, we created a reduced version (AKLIMATE-reduced) that is restricted to the same set of input PID sets as used by BMKL. We refer to the full unrestricted model of AKLIMATE as simply AKLIMATE in this comparison.

AKLIMATE predicted MSI status in the UCEC cohort significantly better than AKLIMATE-reduced or any of the BMKL models (Fig.3a). In particular, AKLIMATE achieved mean AUROC of 0.962 compared to 0.938 for AKLIMATE-reduced (*P <* 4.1*e −* 08; Wilcoxon signed-rank test), suggesting a predictive benefit to AKLIMATE’s larger collection of gene sets. A larger gene set collection is both more likely to contain sets derived specifically to describe the MSI process and more flexible in terms of the possible combinations of component gene sets. Indeed, the most informative feature set according to AKLIMATE was “MSI Colon Cancer” (16.9% relative contribution to model explanatory power) - a gene expression signature for MSI-H vs MSI-L+MSS in COADREAD cohorts [39] (Fig.3b). Futhermore, the next two most informative sets were “GO DNA Binding” (4.55% relative contribution) and “REACTOME Meiotic Recombination” (2.4% relative contribution), both of which are strongly relevant to DNA mismatch repair (MMR). Of note, MLH1 was the top-ranked single feature in AKLIMATE (26.7% relative contribution) and was present in the ten top ranked gene set kernels (Fig.3b). MLH1 is a key MMR gene involved in meiotic cross-over [36] - loss of MLH1 expression, usually through DNA methylation, is known to cause microsatellite instability.

**Fig. 3.**
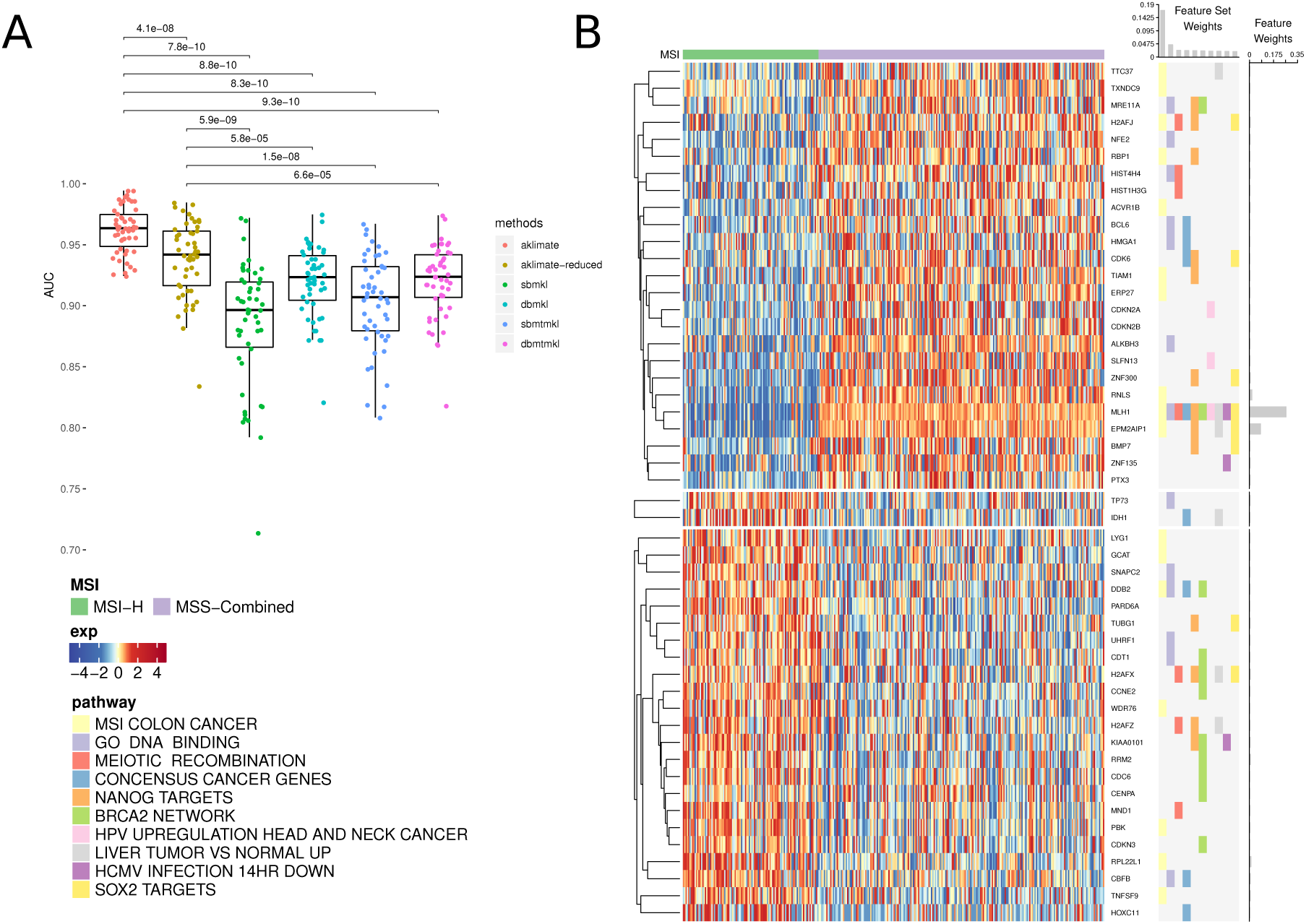
AKLIMATE performance on predicting MSI in UCEC TCGA. A) Performance of AKLIMATE and BMKL on classifying MSI-H vs MSI-L+MSS. AUROC computed for 50 75%/25% stratified train/test splits. P-values for Wilcoxon signed-rank test pairwise comparisons. Methods compared: aklimate - aklimate on full collection of feature sets; aklimate-reduced - aklimate with 196 PID pathways; sbmkl - sparse single-task BMKL; dbmkl - dense single-task BMKL; sbmtmkl - sparse multi-task BMKL; dbmtmkl - dense multi-task BMKL. Multi-task BMKL models are trained to simultaneously predict MSI status on UCEC and COADREAD cohorts. B) Top 10 predictive AKLIMATE feature sets and top 50 predictive features. Expression of top 50 features (left heatmap); Membership of most predictive features in most predictive feature sets (right heatmap). Features are organized by KNN clustering into 3 groups, followed by hierarchical clustering within each cluster. Feature set model weights scaled to sum up to 1 (barplot, top of right heatmap). Feature model weights scaled to sum up to 1 (barplot, right of right heatmap). Feature and feature set weights averaged across 50 train/test splits.

These results demonstrated that AKLIMATE is able to pinpoint individual causal genes as it sifts through thousands of gene sets. In contrast, the meta-pathway constructed by AKLIMATE-reduced represents a poorer approximation to the underlying biological process, as evidenced by its lower AUROC. This is likely due to the fact that, out of the 196 PID pathways used, only “PID P53 Downstream” (35.2% relative contribution) contained MLH1 Fig.S2), limiting the influence of this key gene on the prediction task.

Interestingly, even though AKLIMATE-reduced used the same feature sets as BMKL, it achieved a statistically significant improvement over all BMKL varieties (Fig.3a). This included both the non-sparse and multi-task BMKL versions, despite the fact that the latter benefitted from an entire additional COADREAD data set. In this case, the difference in kernel representations may have contributed to the improved performance of AKLIMATE-reduced (see *Discussion*).

AKLIMATE outperformed other methods on the COADREAD MSI classification task as well. However, the extent of the improvement was smaller because all classifiers perform well on this problem (Fig.S3).

### Breast Cancer Survival

For our second benchmark, we considered the task of predicting survival in the Breast Cancer International Consortium (METABRIC) cohort [19]. This problem is significantly more challenging than predicting MSI status, as demonstrated by the DREAM Breast Cancer Challenge [52] and elsewhere [66]. Another difference between this task and MSI inference is that the METABRIC cohort is annotated with curated clinical data. In fact, the clinical features are quite informative for survival prediction - pre-competition benchmarking by the DREAM Challenge organizers found that models that used exclusively clinical features significantly out-performed ones that used only genomic features, and performed only marginally worse than models in which clinical features were augmented by a subset of molecular features selected through prior domain-specific knowledge [5]. In addition, the best pre-competition clinical feature model had better accuracy than all but the top 5 models in the actual challenge [52].

A breast cancer model that foregoes the use of clinical data would clearly suffer from inferior performance as well as reduced relevance in real-world medical settings. To achieve clinical data integration, AKLIMATE introduces a special category of “global” features, which are added to the “local” features of each feature set prior to the construction of its corresponding RF. Global features can therefore be interpreted as a uniform conditioning step applied to all AKLIMATE component RFs. In our METABRIC analysis, all clinical features are treated as global.

We compared AKLIMATE to two state-of-the art METABRIC survival predictors. The first one, which we refer to as BCC, was the top-performer in the Sage Bionetworks–DREAM Breast Cancer Prognosis Challenge [52][13] - an ensemble of Cox regression, gradient boosting regression, and K-nearest neighbors trained on different combinations of clinical variables and molecular-feature derived metagenes. The second one - Feature Selection MKL (FSMKL)[66] is a pathway-informed extension of SimpleMKL [59] that uses linear and polynomial kernels created from clinical data and molecular features in pathways of the Kyoto Encyclopedia of Genes and Genomes (KEGG) [40]. In FSMKL, pathway features from different data types lead to the construction of separate kernels - in the case of METABRIC, each pathway produces distinct expression and copy number kernels (AKLIMATE, in contrast, learns one kernel matrix from the combined pathway features across all data types). In addition, each clinical feature is treated as a singleton pathway and produces an individual kernel. To make our results directly comparable to FSMKL and BCC as presented in [66], we used a subset of the patient cohort (*N* = 639) and a reduced set of clinical variables to match the dataset used in that publication (see *Supplementary Information*).

We cast the problem as a classification task where algorithms use molecular and clinical data as features to predict whether a patient is alive or not at the 2000 day mark. Based on the 2000 day cutoff, there were 387 survivors and 252 non-survivors in the reduced cohort. Similar to the MSI analysis, we performed 50 stratified repeats of 80% train and 20% test partitions; AKLIMATE was trained on each training split and its accuracy computed on the respective test samples. To decrease computation time, AKLIMATE’s kernel construction step used 1000 trees instead of the default 2000 (see *Supplementary Information* for all other hyperparameter settings).

The full AKLIMATE model had higher mean accuracy than BCC (74.1% vs 73.2%) and was on par with FSMKL (74.1% vs 74.2%). This was unsurprising given that all three models used clinical variables and correctly prioritized them as driver features. Furthermore, two of AKLIMATE’s main advantages were not fully utilized under these experimental settings. First, the benefit of incorporating prior knowledge is reduced because of the relatively low information content of genomic features. Second, the clinical variables may lack complex interactions, which may explain why modeling them with simpler linear techniques is just as effective. In fact, the most explanatory feature in the full AKLIMATE model (16% relative contribution) was the Nottingham Prognostic Index (NPI)[30] - a linear combination of tumor size, tumor grade, and number of lymph nodes involved. Even with these disadvantages, however, AKLIMATE achieved performance on par with two state-of-the-art methods that were specifically optimized for the METABRIC data.

As expected, clinical information proved to be the most influential data type for survival prediction. AKLIMATE models with clinical features alone were significantly more accurate than AKLIMATE models with genomic features alone (p-val=1.9e-08, Fig.4a). This is even more striking considering the models used tens of thousands of genomic features versus only 15 clinical ones. However, the AKLIMATE models using both clinical and genomic features outperformed each single-component model (p-val=1.4e-09 for full versus genomic, p-val=6.7e-04 for full versus clinical, Fig.4a), suggesting that the inclusion of genomic features contained complementary signals with respect to the clinical variables. The mean relative contributions of each data type in the full models were 61.6% for clinical, 29.6% for expression, and 8.8% for copy number.

Of note, while the full AKLIMATE and FSMKL models achieved similar mean accuracy, the importance of individual features and pathways in each model were quite different. AKLIMATE heavily favored clinical variables, with 54.5% of the model’s relative explanatory power carried by just five features - NPI, tumor size, lymph node involvement, age and treatment (Fig.4b). FSMKL ranked tumor size as the most important clinical feature (second overall), with other clinical variables also generally considered relevant - e.g. NPI(9th), age (11th), histological type(14th), tumor group(34th) and PAM50(40th) [66]. Most clinical variables, however, were considered less informative than the top ranked KEGG pathway-based kernels. FSMKL’s highest weighted kernel was “Intestinal Immune Response for IgA production”, followed in importance by “Arachidonic acid metabolism”, “Systemic lupus erythematosus”, “Glycerophospholipid metabolism”, and “Homologous recombination” - all of these KEGG-based kernels scored higher than any clinical variable with the exception of tumor size [66].

**Fig. 4.**
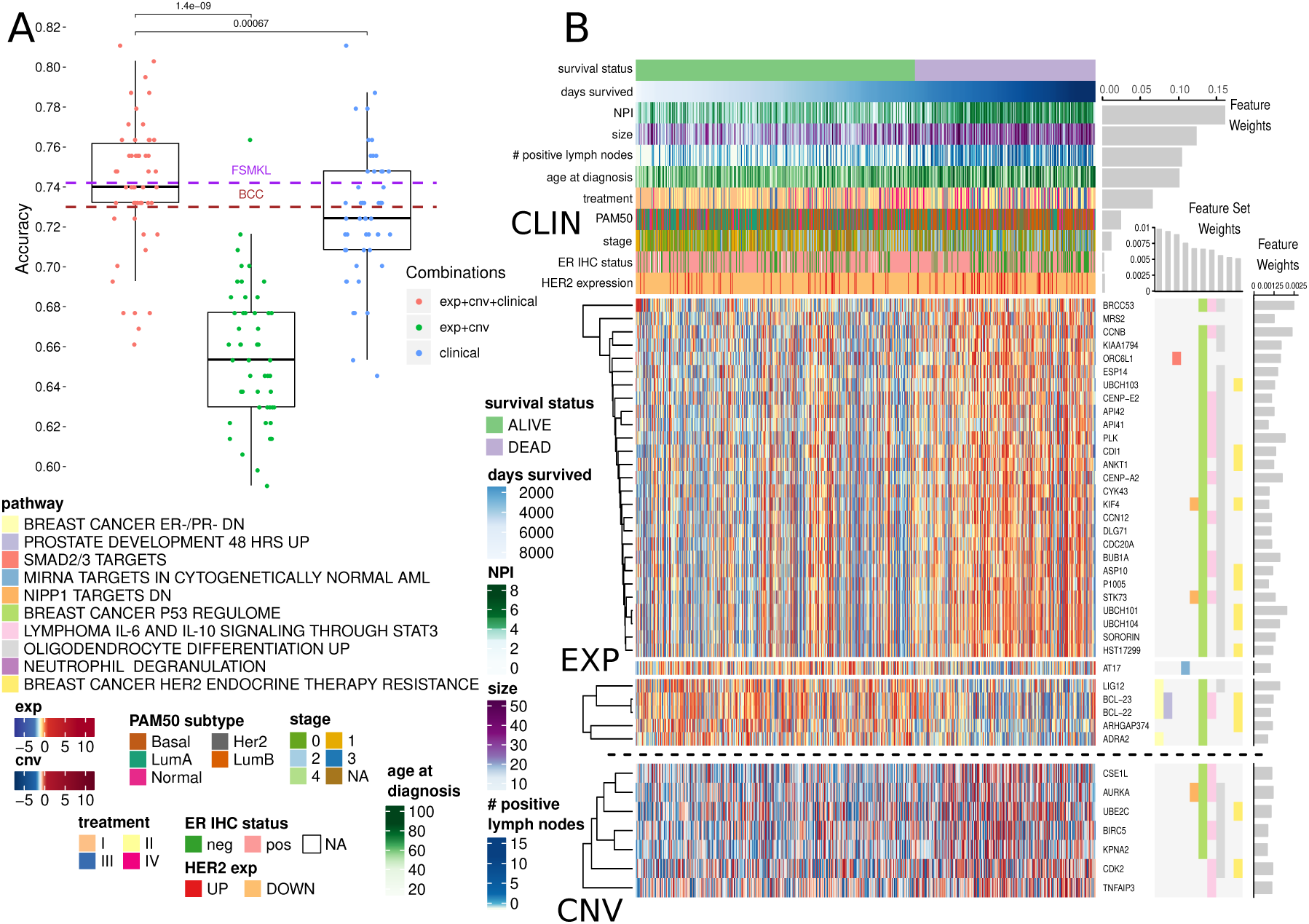
AKLIMATE performance on predicting survival at 2000 days in the METABRIC cohort. A) Performance of AKLIMATE under different data type combinations. EXP+CNV - AKLIMATE with genomic features only; clinical - an RF model run with the clinical variables only; EXP+CNV+CLINICAL - AKLIMATE with genomic features as “local” variables and clinical features as “global” ones. FSMKL and BCC dashed lines show mean performances for the two models under 5-fold cross-validation as shown in [66]. B) AKLIMATE results highlighting the top 10 predictive feature sets and top 50 predictive features. Figure organized as Fig. 3. Clinical variables shown as column annotations; included only if among the 50 most informative features in the model. Clinical variables are ranked from top to bottom by relative predictive contribution. Survival status is a binary variable representing survival at 2000 days (labels) while days survived shows actual duration of survival. Samples sorted by days survived within the two classes. Feature and feature set weights averaged across 50 train/test splits.

While differences in variable importance across methods are expected, further analysis is necessary to determine which model best aligns with the relevant biology of patient outcomes. Encouragingly, AKLIMATE’s most informative feature sets were enriched for breast cancer progression and response to treatment, with 3 of the top 10 and 8 of the top 20 (Table S1) related to these functional groups. For example, the most informative feature set (“BREAST CANCER ER-/PR- DN”) represents a signature that is correlated with reduced protein abundance of the estrogen (ER) and progesterone (PR) hormone receptors [17]. Similarly, the 9th most informative pathway (“BREAST CANCER HER2 ENDOCRINE THERAPY RESISTANCE”) [16] captures transcriptome changes associated with the development of resistance to targeted therapies. Both of these signatures serve as proxies for highly relevant information not available for the METABRIC cohort (protein activity for PR and therapy resistance). Furthermore, the latest AJCC breast cancer staging manual introduces tumor grade and ER/PR/HER2 receptor status among the key breast cancer biomarkers, which already include tumor size and lymph node engagement [26]. All of these appeared as highly informative AKLIMATE model features, either directly or via a proxy genomic signature.

### shRNA knockdown viability

We used the problem of predicting cell viability post shRNA knockdown to showcase AKLIMATE’s ability to integrate multiple data types and solve regression tasks. In this case, the prediction labels were continuous values representing cell line survival after the shRNA-mediated mRNA degradation of a particular gene. Gene profiles were computed with ATARIS [67] - they are consensus scores that combine viability phenotypes from multiple shRNAs targeting the same gene. We selected 37 such profiles for different genes across 216 cancer cell lines from the Cancer Cell Line Encyclopedia (CCLE) [15]. We chose these 37 tasks (out of 5711 available consensus profiles) because they had at least 10 cell lines showing strong (*>*2 sd from mean) knockdown viability response and are in the top quartile by variance of all consensus profiles (see *Supplementary Information*). Based on the results of the DREAM9 gene essentiality prediction challenge [29], we expected this task to be significantly harder than the other case studies.

We used expression and copy number measurements from CCLE [4] as predictive features. We augmented these two data types by adding discrete copy number alteration calls made by GISTIC2 [53] and activities for 447 transcriptional and post-transcriptional regulators inferred by hierarchical VIPER [78][61]. We focused on the 206 cell lines for which we have knockdown profiles, copy number and expression features.

We compared AKLIMATE’s performance to three of the five top performing methods in DREAM9 [29] as well as three baseline algorithms. We briefly describe the DREAM9 sub- challenge 1 top performers next. Multiple Pathway Learning (MPL)[78] took 5th place (see https://www.synapse.org/#!Synapse:syn2384331/wiki/64760 for challenge results) - it used elastic-net regularized Multiple Kernel Learning with Gaussian kernels based on feature sets from the Molecular Signature Database (MSIGDB) [47]. MPL and Random Forest Ensemble (MPL-RF) took 2nd place - its prediction was based on averaging a Random Forest classifier with MPL. MPL and MPL-RF were our contributions to the DREAM9 challenge. Both methods were run with the same hyperparameters and pathway collections as in DREAM9 [29][78]. Kernelized Gaussian Process Regression (GPR) took 3rd place - it used extensive filtering steps to reduce the input feature dimensionality, followed by principal component analysis, and finally Gaussian Process regression with covariance computed from the principal components [29] (see also https://www.synapse.org/#!Synapse:syn2664852/wiki/68499 for implementation and model description). We downloaded the code from the Synapse URL and ran it with the published DREAM9 hyperparameters.

To provide performance baselines, the DREAM9 winners were augmented by standalone Random Forest (RF), Generalized Linear Model (GLM) with lasso penalty (GLM-sparse), and GLM with L2 regularization (GLM-dense). RF was run with the *ranger* R package [85] with the following hyperparameters - sampling without replacement with 70% of the samples used for tree construction, minimum node size of 10, 1500 trees and 10% of the features randomly sampled for each node split. GLM-dense and GLM-sparse were run using the *glmnet* R package [24] with the response family set to “gaussian” and the strength of regularization *λ* optimized through cross-validation. The elastic net tradeoff between the lasso and ridge penalties *α* was set to *α* = 0.8 (GLM-sparse) and *α* = 0.001 (GLM-dense).

We did not include the nominal first place winner of DREAM9 - an ensemble of four kernel ridge regression models with kernels trained through Kernel Canonical Correlation Analysis and Kernel Target Alignment - because we could not re-run the source code supplied with the challenge submission. We feel this omission is not material as the top 3 methods were declared joint co-winners - their results were shown to be statistically indistinguishable from each other but separable from the rest of the entries [29]. Furthermore, experiments in [78] suggest that this method underperforms MPL, MPL-RF and GPR when only high-quality shRNA knockdown profiles are considered.

To save computational time, we compared methods on a single stratified train/test split for each ATARIS profile (a different split for each profile) where 67% of the cell lines were used for training and 33% were withheld for testing. Each method was run with its recommended parameters and filtering steps - if no filtering steps were specified, the method used all available features. AKLIMATE’s prediction binarization quantile was set to its default value of *q* = 0.05 (see *Methods*, also *Supplementary Information* for all other settings).

AKLIMATE achieved average Root Mean Squared Error (RMSE) of 1.031 vs 1.047 for GPR, 1.055 for RF, 1.065 for GLM-dense, 1.07 for MPL, 1.071 for GLM-sparse and 1.08 for MPL-RF. The mean RMSE difference was statistically significant in all but one case (Fig.5A, Wilcoxon signed rank test). AKLIMATE was also the top performer when we considered the number of times an algorithm achieved the best RMSE on an individual prediction task (Fig.5B, AKLIMATE retained top spot under non-RMSE metrics as well - Fig.S4). AKLIMATE performed better than average across nearly all tasks (Fig.S5); its advantage was particularly pronounced in predicting the essentiality of key regulators (CTNNB1, FOXA1, MDM4, PIK3CA) or housekeeping genes (PSMC2, PSMC5). As these gene classes are heavily studied and thus over-represented in our pathway compendium, AKLIMATE’s enhanced accuracy may be due to the relatively higher abundance of relevant prior knowledge.

For example, AKLIMATE’s ability to predict MDM4 knockdowns benefited from MDM4’s function as a p53 inhibitor - many of the most informative feature sets in the MDM4 model relate to p53’s regulome and its role in controlling apoptosis, hypoxia and DNA damage control (Fig.5C). The top features were also functionally linked to p53 - CDKN1A (28.6% relative contribution) is a kinase that controls G1 cell cycle progession and is tightly regulated by p53; MDM2 (8% relative contribution) participates in a regulatory feedback loop with p53; ZMAT3 (3.7% relative contribution of expression; 3.5% of copy number) is a zinc finger whose interaction with p53 plays a key role in p53-dependent growth control. In addition, the four p53 features from each data type were all present among the 50 most informative ones. Individually they carried little signal (relative contribution: 0.5% expression, 0.3% copy number, 0.3% inferred protein activity, 0.2% GISTIC score) but taken together they clearly implicated p53 as one of the top 10 most informative genes. The ability to capture such multi-omic interactions is one of AKLIMATE’s main strengths, made possible by the use of RF kernels that fully integrate all data types. The synergy between different p53-based features is impossible to observe in methods that assign individual kernels to each data type (e.g. BMKL, FSMKL and MPL). Encouragingly, AKLIMATE dominated MPL/MPL-RF on almost all tasks even though they share the same MKL solver (Fig.5A, Fig.S5).

FOXA1 knockdown was another example of AKLIMATE showing superior predictive performance on a well-studied gene. FOXA1 dysregulation is an essential event in breast cancer progression and subtype characterization. Due to breast cancer’s prevalence and clinical importance, there is an extensive list of relevant signatures in our feature set compendium. Eight of the top 10 (and 12 of 14 overall) feature sets in the FOXA1 AKLIMATE model were directly related to breast cancer experiments under different conditions (Fig. S7). As expected, FOXA1 features were the most informative (relative contribution: 31.5% FOXA1 expression; 3.4% FOXA1 inferred activity), with AR also among the top 5 most informative genes (relative contribution: 2.8% AR expression).

Out of the 37 shRNA prediction tasks we considered, KRAS was the most obvious example of a well-characterized gene that did not experience discernible accuracy improvement over competing methods (Fig.S5). Our hypothesis is that a true biological “driver” is absent from the set of molecular features presented to AKLIMATE. To test this, we added a mutation data type containing information for 8 key regulators (KRAS, NRAS, PIK3CA, BRAF, PTEN, APC, CTNNB1 and EGFR), 3 of which (KRAS, CTNNB1, PIK3CA) have knockdown profiles among the 37 shRNA prediction tasks.

Even though our mutation data type consisted of only 8 features, its addition led to a dramatic improvement in KRAS shRNA prediction accuracy across all metrics (Fig.5D, mean RMSE 0.918*±*0.025 vs 1.02*±*0.024; mean Pearson 0.665*±*0.023 vs 0.529*±*0.025; mean Spearman 0.57*±*0.039 vs 0.453*±*0.034). Furthermore, the KRAS mutation feature was by far the most informative (23.5% relative contribution, Fig.5E), followed by KRAS copy number (6.9%), KRAS GISTIC (3.3%) and KRAS expression (3.1%). The addition of the KRAS mutation feature was not only key to improving the predictive performance of the model, but also helped prioritize KRAS features from other data types. While KRAS expression, copy number and GISTIC features all appeared among the 50 most informative ones in the “no mutation” run, their combined relative contribution was only 3.33% (1.5% copy number, 1% expression, 0.8% GISTIC, Fig.S8). This provides another example of the ability of AKLIMATE’s RF kernels to highlight composite patterns of multi-omic interactions.

Our training features are measured at the gene level, but AKLIMATE has no constraints on the inclusion of more granular data. For example, it can evaluate the importance of mutations at particular amino acid positions or the type of substitutions caused. In the case of KRAS, glycine replacement by either aspartic acid or valine in the 12th amino acid position appears to have the biggest negative effect on cell viability post-shRNA knockdown (Fig.5E). G12 is a well-known KRAS mutation hotspot [34] - AKLIMATE’s ability to prioritize relevant hotspots can be a key advantage in modeling drug response or recommending treatment strategies.

The addition of mutation data did not yield any predictive benefit in modeling PIK3CA or CTNNB1 post-knockdown viability (Fig. S9-S10). CTNNB1 had only 9 mutations in the cohort, all of them in different codons. PIK3CA had 30 mutations, with some hotspots - the lack of improvement in this case may be due to the change in protein sequence not being biologically relevant, the ability of other genomic features to fully capture the mutation signal, or the fact that the “no mutation” models were quite accurate to begin with.

## Discussion

Recent surveys of cancer genome landscapes have shown that alterations of a particular pathway can involve many different genes and many different kinds of disruptions - for example, RB1 mutation, RB1 methylation, or CDKN2A deletion could all lead to aberrant cell proliferation [21]. Consequently, many bioinformatics approaches seek to combine data at the level of a biological process to benefit machine-learning applications in the cancer genomics setting. However, data platform diversity often prohibits such integration - variables can be of different scales (e.g. copy number vs gene expression) or different types (continuous DNA methylation, binary mutation calls, ordinal inferred copy number estimates). AKLIMATE’s early integration approach is a potential solution to capturing complementary process-level information spread across data modalities - all data types are considered, and potentially used, when constructing a process-related RF kernel. AKLIMATE does so by building supervised tree-based empirical kernel functions that optimally align the training labels with each process-specific set of multimodal data. In contrast, MPL, FSMKL and other MKL approaches with unsupervised kernel construction compute segregated data-type specific kernels for each pathway and let the linear combination “meta-kernel” determine their optimal combination. This may result in suboptimal solutions - features that belong to the same pathway but are in different data types can now only interact with each other on the kernel level and not individually. AKLIMATE’s approach creates a richer interaction model that is flexible enough to capture same-gene, cross-gene, and cross-data type interactions.

Another limitation of current pathway-informed kernel learning methods is that a single informative feature can go undetected if only present in large pathways - if all member features contribute equally to kernel construction, the importance of the relevant feature is obscured by the non-relevant majority. In contrast, AKLIMATE’s RF kernels effectively allow individual features to influence the model. This advantage is illustrated by the improvement in AKLIMATE performance over BMKL on the MSI prediction task. BMKL’s Gaussian kernels treat each feature in a feature set as equally important in the computation of the respective kernel matrix. As MLH1 appears only once in the PID pathway compendium, its contribution is masked by less informative features. On the other hand, due to the supervised manner of their construction, AKLIMATE’s RF kernels inherit an RF’s ability to prioritize features based on their relevance to the classification task - informative features are by definition overrepresented among tree node splitting variables. As all components of the RF kernel are derived from properties of the RF trees, informative features exercise proportionately higher influence over the RF kernel construction. Therefore, if only a small subset of a pathway’s features are truly relevant, they can be clearly distinguished from (thousands of) non-relevant ones. For example, both AKLIMATE and AKLIMATE-reduced pick MLH1 as the most informative feature (26.7% and 16.4% relative contribution respectively), with a steep decline in the importance of the next best feature (AKLIMATE - EPM2AIP1, 8% relative contribution; AKLIMATE-reduced - PARD6A, 4.8% relative contribution).

A further key advantage of AKLIMATE is its ability to accommodate variables that do not readily map to genome-based feature sets (e.g. clinical data). In cases such as METABRIC, where clinical features provide much of the predictive power, the most salient question becomes what genomic features can add orthogonal information given that the clinical data are already incorporated into the model. Posing the problem as a conditional relationship between clinical and molecular data closely reflects the practical situation in hospital settings where clinical information is nearly always available to treating physicians. AKLIMATE and FSMKL illustrate two different ways of incorporating such information. FSMKL treats each clinical variable as a feature set of size one and constructs a kernel for each of them individually. This approach is viable, but quite restrictive in how it models clinical variable interactions. While AKLIMATE can accommodate such a setup, it also permits a more complex representation of the way features interact - each clinical feature is of a special “global” type that gets included in every feature set in addition to the features “local” to it. The “global”-”local” feature hierarchy allows maximum flexibility in modeling interactions among clinical variables and between clinical variables and genomic features. Such a hierarchy is necessary when features work on different biological scales - for example, tumor grade is an organ-level characteristic that captures a snapshot of the behavior of millions of cells and is therefore likely to be vastly more informative than the copy number status of an individual gene.

An important aspect of AKLIMATE’s use of prior knowledge is its ability to identify relevant features even in cases where many confounders exhibit high collinearity. This problem is similar to the one encountered in genome-wide association studies where an allele conferring a phenotype of interest can exist in a large haplotype block containing the alleles of many other irrelevant “hitchhiking” genes. In such situations, prior knowledge often helps researchers select the true causal variant among potentially many false positives. Consider AKLIMATE’s top two most informative features for the MSI prediction task – MLH1 and EPM2AIP1. EPM2AIP1 is on the DNA strand opposite to MLH1, shares a bi-directional CpG island promoter with it and can be concurrently transcribed [31]. The transcriptional profiles of the two genes are nearly identical (Fig.3b) - in the absence of other information it would be extremely difficult to prioritize the “driver” (MLH1) over the “passenger” (EPM2AIP1) using expression data alone. AKLIMATE’s feature sets provide the necessary prior knowledge - while EPM2AIP1 is indeed deemed the second most informative feature, its relative contribution is over three times smaller than that of MLH1.

AKLIMATE’s robustness to false positives is not limited to expression features - it can prioritize relevant genes even if they are subject to large-scale copy number events and thus have almost identical copy number profiles with many other genes. We observed this effect in the MDM4 knockdown prediction task (Fig. 5C) as well as KRAS knockdown prediction with mutation data (Fig. 5E). In the former, there is a clear large scale copy number event that involves 7 of the 50 most predictive genes, but ZMAT3 is given by far the highest weight because of the biological prior of the feature sets. Similarly, in the latter 9 copy number features have very similar profiles, but the KRAS one is prioritized as the most important. The KRAS GISTIC feature is also favored among a group of genes affected by large-scale events - an indicator that the robustness to collinearity extends beyond continuous to categorical features.

**Fig. 5.**
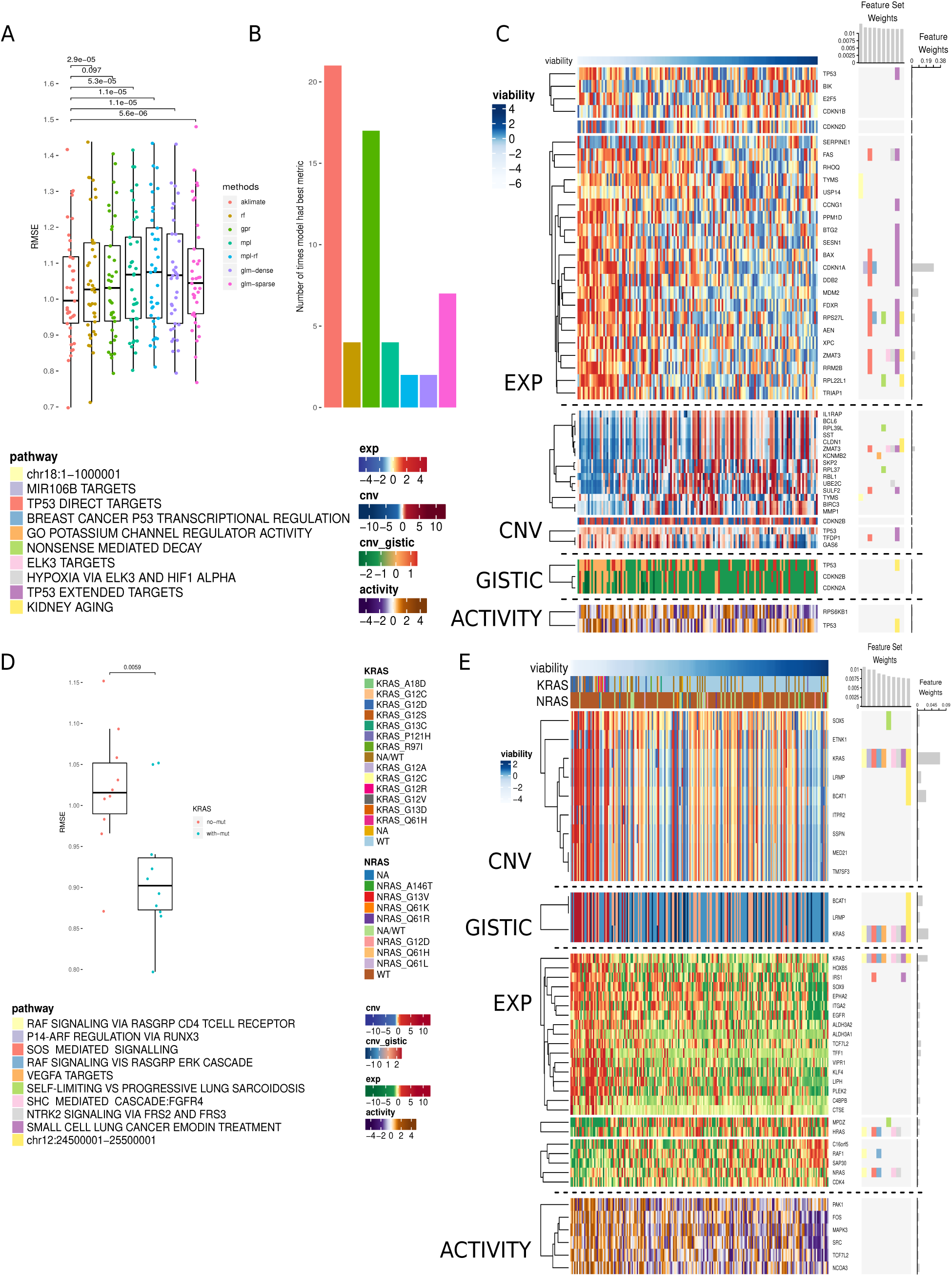
Prediction of cell line viability after shRNA gene knockdowns. A) RMSEs of AKLIMATE and competing methods on 37 consensus viability profiles from the Achilles dataset. Methods: Random Forest (RF), Gaussian Process Regression (GPR), Multiple Pathway Learning (MPL), ensemble of MPL and Random Forest (MPL-RF), L2 regularized linear regression (GLM-dense), L1 regularized linear regression (GLM-sparse). B) Number of times an algortihm produced the best RMSE on a prediction task. To prevent small relative RMSE differences from having a biasing effect on the win counts, for each task we consider all algorithms with RMSE within 1% of the min RMSE to be joint winners. For that reason total win counts add up to more than the number of regression tasks. C) AKLIMATE top 10 informative feature sets and top 50 informative features for the task of predicting MDM4 shRNA knockdown viability. Organized as Fig. 3. D) RMSEs of KRAS AKLIMATE models with and without the use of mutation profiles for 8 key regulators. Results shown for 10 matched stratified train/test splits where 80% of the cohort is used for training and 20% for testing. E) AKLIMATE top 10 informative feature sets and top 50 informative features for the task of predicting KRAS shRNA knockdown viability when mutation features are used (feature and feature set weights averaged over 10 train/test splits). Organized as Fig. 3.

Making use of prior knowledge, such as the results of past experiments, is often a key component in the success of machine-learning applications in genomics analysis [5] [75]. AKLIMATE uses a biologically-motivated prior distribution on the feature space - as demonstrated, this approach often outperforms methods that use a uniform prior over input features. AKLIMATE updates its prior information in a data driven manner - therefore, the feature set compendium does not have to be tailored to the problem at hand, avoiding the need for problem-specific filtering heuristics. When high quality relevant prior experiments are available, AKLIMATE tends to perform better. For example, the most important feature set for MSI prediction in endometrial cancer is a previously published signature characterizing MSI in colon cancer (Fig.3B). From that perspective, AKLIMATE acts as a framework for prioritizing past experiments that are most relevant to the interpretation of a new dataset.

By aggregating feature weights within individual data types, AKLIMATE can also be used to rank data type contributions. For example, while expression is generally most important in predicting shRNA knockdowns (mean importance 71.6% across tasks), there are cases where copy number is more informative, such as PSMC2(61.7%), RPAP1(55%) and CASP8AP2(47%) (Fig. S6). Furthermore, inferred protein activity varies tremendously in terms of its contribution - from *<* 1% in predicting PSMC2 knockdowns to 27.7% for STRN4 (Fig. S6). AKLIMATE’s ability to zero in on information-rich data types could help in designing targeted future experiments.

From a stacked learning perspective, AKLIMATE augments standard level-one cross validated predictions with topological aspects of the base RFs - how often data points end up in the same leaf node and whether they tend to end up in early-split or late-split leaves. Such augmentation improves performance - in all case studies AKLIMATE outperforms both the best individual base learner and the ensemble that averages base learner predictions (Fig. S1). In addition, AKLIMATE does better than a standard Super Learner (regularized linear regression meta-learner) although without achieving statistical significance on some of the data sets (Fig. S1). This suggests that propagating additional information from a base learner (beyond the predictions it makes) into the level-one data can lead to a more accurate meta-learner. While AKLIMATE provides a blueprint for tree-based algorithms, using other base learners might yield even better results.

AKLIMATE requires minimal feature pre-processing, can query tens of thousands of feature sets and its main steps are trivially parallelizable. It performs as well as, or better than, state-of-the-art algorithms in a variety of prediction tasks. AKLIMATE can natively handle continuous, binary, categorical, ordinal and count data, and its feature sets are easily extendable. For example, gene expression data can be augmented by epigenetic measurements (e.g. DNA methylation or ATAC-Seq), mutations in promoters or enhancers, alternative splice variant proportions, and so on. Furthermore, slight modifications to AKLIMATE could incorporate prior importance scores for features within a feature set as well as structured relationships such as known feature-feature interactions (e.g. transcription factor to target gene). To simplify the exposition, we have focused our derivations and applications to regression and binary classification problems. However, AKLIMATE is readily extendable to the multi-class setting and the current public code base provides such an implementation.

While AKLIMATE demonstrates enhanced predictive power, future improvements are certainly possible that could lead to a bigger performance boost. For example, a kernel that captures the topological distance between samples in RF trees as suggested in [22] could provide a more accurate replacement for the *K*_2_ kernel we introduced. Sample-weighting schemes during kernel construction could be investigated as well. For example, [11] suggests a weighted RF kernel where sample contributions depend on the predictive accuracy of their asigned leaves or the classification error rate for an individual sample.

Finally, even though we present case studies from the field of bioinformatics, AKLIMATE can be applied to any task with multi-modal data and prior knowledge in the form of feature groups, particularly when the feature groups have evidence that span data types.

## Supplementary Information

### Data Acquisition

#### Microsatellite Instability

Data for the COADREAD and UCEC TCGA cohorts were downloaded from the Synapse copy of the PANCAN12 TCGA cohort [32] (synapse object id syn300013, https://www.synapse.org/#!Synapse:syn300013/wiki/70804). The UCEC expression data (syn1446289) were log transformed. The log-transformed data (20,501 features) were used as the basis for BMKL- (see [27]) and AKLIMATE-specific filtering (see *AKLIMATE pre-processing*). The AKLIMATE filtered data set contained 13,424 expression features. The MSI status for UCEC patients was extracted from UCEC clinical data (syn1446167).

Similarly, the COADREAD expression dataset was created by joining the COAD (syn1446197) and READ (syn1446276) PANCAN12 cohorts, log transforming the combined matrix and applying respective filtering steps. The AKLIMATE filtered data set contained 14,036 features. MSI status was downloaded from `firebrowse.org` and matched to the whitelisted samples for the joint COADREAD PANCAN12 expression set.

#### METABRIC Survival

Expression (Illumina HT12 array), copy number (Affymetrix SNP 6.0) and clinical data for the METABRIC cohort [19] were downloaded from https://www.synapse.org (Synapse ID syn1688369). Expression and copy number data were processed as described in the marker paper [19]. Expression data used Illumina HT12V3 probe identifiers while copy number data had Entrez gene features. Since our pathway compendium was HGNC-based, we translated each HGNC gene set to all Illumina probes and Entrez gene IDs matching any of its members. We used the *IlluminaHumanv4*.*db* Bioconductor package for the Illumina-HGNC map and the *org*.*Hs*.*eg*.*db* package for the Entrez-HGNC map.

To match the analysis in [66], we restricted the METABRIC cohort to 639 patients (list obtained in personal communication with authors). For the same reason, we did not use the full set of clinical information available, but limited it to variables used in [66], namely:

1. Age at diagnosis
2. Tumor size
3. Tumor grade
4. Tumor stage
5. Number of positive lymph nodes
6. Histological type
7. Estrogen receptor IHC status and expression-based status
8. HER2 IHC status, SNP6 status, and expression-based status
9. Nottingham Prognostic Index
10. PAM50-based breast cancer subtype
11. Cellularity
12. Composite treatment status

The last clinical variable was not present in the METABRIC clinical file, but was created by integrating aspects of other clinical variables in a manner described in [66].

Expression and copy number data sets were filtered as described in *AKLIMATE pre-processing*, with mean and variance calculations based on the reduced rather than the full cohort. The combined AKLIMATE filtered feature set contained 20,022 expression, 8608 copy number and 15 clinical features.

Accuracies for FSMKL and BCC methods were taken from [66].

#### shRNA Knockdown Profiles

Achilles 2.4.3 shRNA knockdown profiles were downloaded from https://depmap.org. The data release contained ATARIS [67] gene-level profiles for 5711 genes across 216 CCLE cell lines. Each profile was computed by aggregating the profiles of multiple shRNAs targeting an individual gene. ATARIS was run with a threshold of *p* = 0.05 on the samples and shRNAs that passed QC inspection (see online QC manifest of the Achilles 2.4.3 data). The mutation profiles of 8 regulators (KRAS, NRAS, PIK3CA, BRAF, PTEN, APC, CTNNB1 and EGFR) were extracted from the sample annotation file for the Achilles 2.4.3 data release.

Matching expression and copy number characterizations of individual cell lines were downloaded from the Cancer Cell Line Encyclopedia (CCLE, https://portals.broadinstitute.org/ccle/data). Expression was measured using Affymetrix U133 Plus 2.0 array, aggregated via Robust Multi-array Average and quantile normalized (see 2012 expression data release on CCLE website). Copy number was evaluated with Affymetrix SNP 6.0 arrays and segmentation of the normalized log2 probe ratios via Circular Binary Segmentation (see 2012 copy number release on CCLE website). GISTIC2 version 2.0.22 was run on the copy number data with default parameters (amplification and deletion thresholds of 0.1, broad event threshold of 0.7, enabled arm level peel-off events, and gene collapsing set to “extreme”). Hierarchical VIPER (hVIPER) was run on the expression data as described in [78] and [61].

Expression (18,900 features), copy number (23,316 features), GISTIC (24,924 features) and hVIPER activity (447 features) data types were provided to all methods for method-specific preprocessing (see main text). In case no pre-processing was specified, a method used all available features.

AKLIMATE used the filtering steps described in *AKLIMATE pre-processing*. The combined AKLIMATE filtered feature set contained 13,652 expression, 10,086 copy number, 9,557 GISTIC and 447 hVIPER activity features.

#### Feature Sets

We used four main sources for biologically relevant feature sets:

1. C2 (Curated Gene Sets) and C5 (Go Gene Sets) collections of MSigDB [72].
2. The GeneSigDB curated collection of published signatures [18].
3. Pathway Commons [12]- a database of databases covering the spectrum of metabolic, molecular, signaling, regulatory and genetic interactions.
4. Gene sets related to chromosomal location. These were constructed by passing TCGA LIHC segmented copy number data through the *CNRegions* function of the *iClusterPlus* [54] R package, with *ϵ* = 0.0025.

Gene sets were excluded from the Small Molecular Pathway Database [38] (SMPDB, part of Pathway Commons) due to the high redundancy and small size of many of the signatures. The “canonical” pathways C2 CP of the MSigDB subcollection were removed due to their high degree of overlap with pathways contained in the more extensive Pathway Commons resource. Finally, all sets with more that 1000 members were removed to maintain specificity. The final compendium consisted of 17,273 gene sets with median size of 30 (min size of 1, max size of 991).

#### AKLIMATE pre-processing

Data for all three case studies was processed in the following manner:

1. Expression data was filtered based on the mean and variance of genes across samples - any gene whose mean or variance falls in the bottom 20% of the mean/variance empirical distribution was discarded.
2. Similar to expression, copy number data, if available, was filtered using a cutoff set at 50%. Whenever GISTIC2 [53] discretized gene-level copy number calls were used they were first filtered in the same manner.
3. The 447 protein activity scores from hVIPER [78][61] representing transcription factor and kinase regulator features were not filtered.

To speed up computation in the larger cohorts, expression and copy number filtered data for the two classification tasks (MSI and METABRIC survival prediction) were discretized by computing the quintiles of the distribution of each molecular feature and binning each quintile into a separate category. Finally, unordered categorical features (e.g. METABRIC clinical variables) were one-hot encoded prior to use by AKLIMATE.

#### AKLIMATE hyperparameters

AKLIMATE was run with the same gene set collection across all prediction tasks (see *Feature Sets*). To increase robustness, gene sets that had fewer than 15 features across all data modalities considered were discarded. Since different case studies use a different number of data types, this thresholding causes the number of eligible gene sets to be task-specific.

The same AKLIMATE hyperparameters were used across all case studies, except for minor deviations described in the main text. To reduce computation time, AKLIMATE component RFs were trained with 50% sampling without replacement - i.e. each tree was grown on a randomly selected 50% subsample of the training set. Studies have shown that this setup performs as well as bootstrapping in predictive accuracy benchmarks [23]. In addition, the trees in each RF base model were set to have minimum leaf size equal to 1% of the size of the training cohort. For the selection of the best RF models **Δ**\*, each RF contained 500 trees. Prior to kernel construction, the forests of **Δ**\* are re-grown with 2000 trees each. Higher number of trees and smaller leaf size tend to provide better approximation to the class discrimination boundary, as demonstrated in [10].

It is generally recommended to keep *mtry* low- e.g. 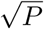 (*P* - total number of features) for classification and 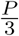 for regression problems because decorrelation among the predictions of ensemble components (e.g. a RF) often leads to an improved performance of the ensemble (see *Methods*). AKLIMATE, however, is an ensemble of ensembles - decorrelation can also be achieved by selecting component RFs that describe independent gene sets. As a consequence, we could prioritize bias reduction within individual RFs - we recommend *mtry* values in the 25-75% range. In our experience, a setting of 25% is fast and accurate. Thus, the number of features randomly selected to try at each node (*mtry*) was set to 25% of the size of the queried gene set.

Finally, we used two different importance metrics for the calculation of feature and feature set relevance - Actual Impurity Reduction (AIR) [55] for classification tasks (microsatellite instability and METABRIC survival), and permutation analysis [8] for regression problems (shRNA knockdown viability). See *Methods* for more details.

#### Implementation

We used the R package *ranger* [85] for calculations involving AKLIMATE’s base RF learners, including permutation-/AIR-based variable importance. We chose *ranger* because of its flexibility in handling splitting rules, variable importance approaches, and learning tasks. It is also one of the fastest RF algorithms currently available, particularly in problems where the number of features is much larger than the number of data points.

For our MKL learner, we ported *SpicyMKL* [74] to R. We chose *SpicyMKL* because its guaranteed super-linear convergence makes it possible to handle thousands of kernels. Furthermore, *SpicyMKL*’s elastic-net regularization allows maximum flexibility in terms of the number of kernels included in the optimal solution. Our R implementation of *SpicyMKL* called *SPICER* is available at https://github.com/VladoUzunangelov/SPICER.

Finally, an R implementation of AKLIMATE is available at https://github.com/VladoUzunangelov/aklimate.

## Supplementary Tables

**Table S1.**
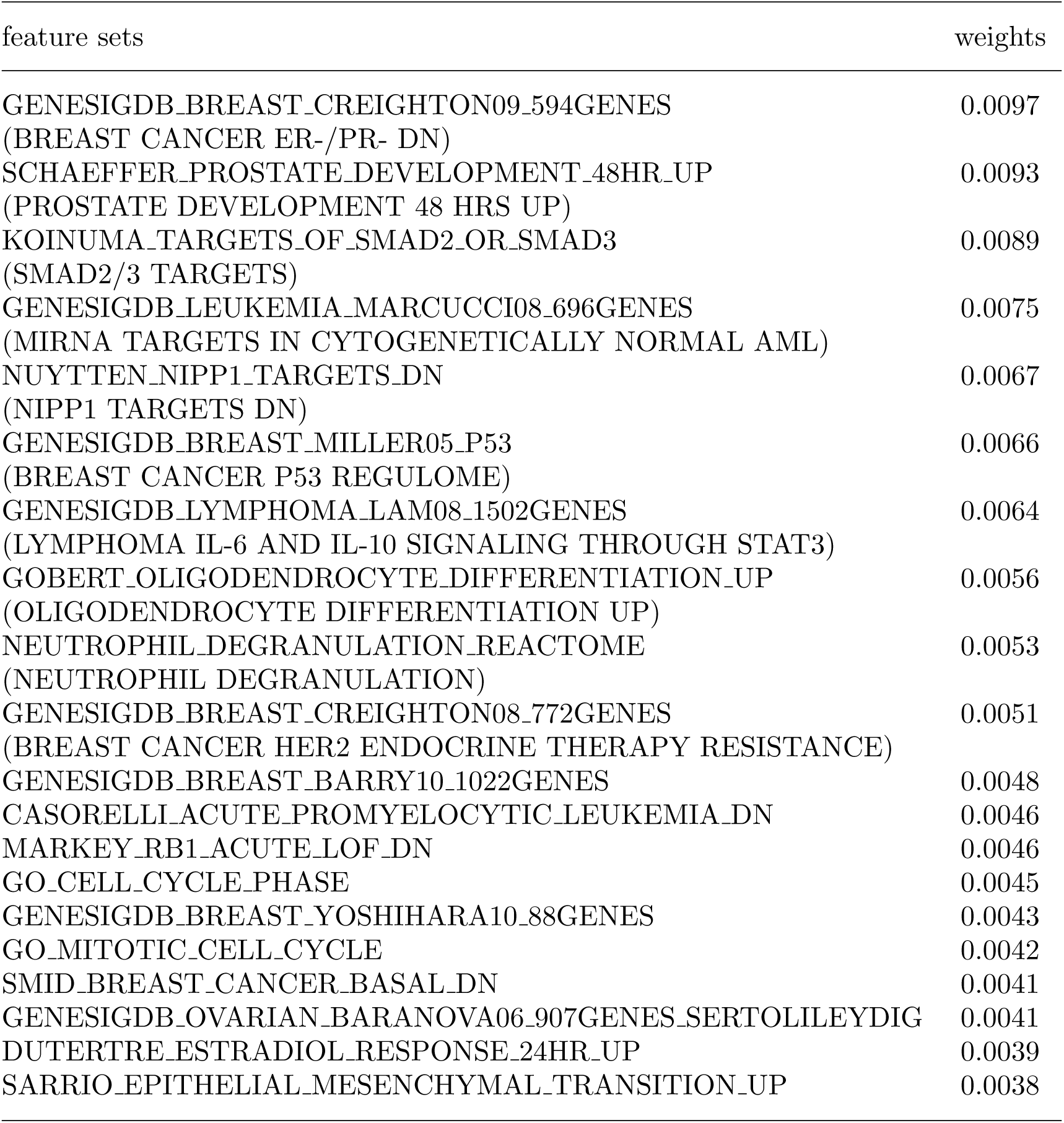
Most informative feature sets for breast cancer survival prediction in METABRIC data. AKLIMATE weights were averaged over 50 train/test splits. The table lists the 20 most relevant feature sets, out of 1836 feature sets with a non-zero weight in at least one train/test split. Weights were normalized to sum to 1. Aliases for the top 10 feature sets are included in brackets - they provide a more descriptive name for the underlying biological function, based on information gathered from the source publication. The aliases are used in Fig. 4 and throughout the text.

| feature sets | weights |
| --- | --- |
| GENESIGDB_BREAST_CREIGHTON09_594GENES<br>(BREAST CANCER ER-/PR- DN) | 0.0097 |
| SCHAEFFER_PROSTATE_DEVELOPMENT_48HR_UP<br>(PROSTATE DEVELOPMENT 48 HRS UP) | 0.0093 |
| KOINUMA_TARGETS_OF_SMAD2_OR_SMAD3<br>(SMAD2/3 TARGETS) | 0.0089 |
| GENESIGDB_LEUKEMIA_MARCUCCI08_696GENES<br>(MIRNA TARGETS IN CYTOGENETICALLY NORMAL AML) | 0.0075 |
| NUYTTEN_NIPP1_TARGETS_DN<br>(NIPP1 TARGETS DN) | 0.0067 |
| GENESIGDB_BREAST_MILLER05_P53<br>(BREAST CANCER P53 REGULOME) | 0.0066 |
| GENESIGDB_LYMPHOMA_LAM08_1502GENES<br>(LYMPHOMA IL-6 AND IL-10 SIGNALING THROUGH STAT3) | 0.0064 |
| GOBERT_OLIGODENDROCYTE_DIFFERENTIATION_UP<br>(OLIGODENDROCYTE DIFFERENTIATION UP) | 0.0056 |
| NEUTROPHIL_DEGRANULATION_REACTOME<br>(NEUTROPHIL DEGRANULATION) | 0.0053 |
| GENESIGDB_BREAST_CREIGHTON08_772GENES<br>(BREAST CANCER HER2 ENDOCRINE THERAPY RESISTANCE) | 0.0051 |
| GENESIGDB_BREAST_BARRY10_1022GENES | 0.0048 |
| CASORELLI_ACUTE_PROMYELOCYTIC_LEUKEMIA_DN | 0.0046 |
| MARKEY_RB1_ACUTE_LOF_DN | 0.0046 |
| GO_CELL_CYCLE_PHASE | 0.0045 |
| GENESIGDB_BREAST_YOSHIHARA10_88GENES | 0.0043 |
| GO_MITOTIC_CELL_CYCLE | 0.0042 |
| SMID_BREAST_CANCER_BASAL_DN | 0.0041 |
| GENESIGDB_OVARIAN_BARANOVA06_907GENES_SERTOLILEYDIG | 0.0041 |
| DUTERTRE_ESTRADIOL_RESPONSE_24HR_UP | 0.0039 |
| SARRIO_EPITHELIAL_MESENCHYMAL_TRANSITION_UP | 0.0038 |

## Supplementary Figures

**Fig. S1.**
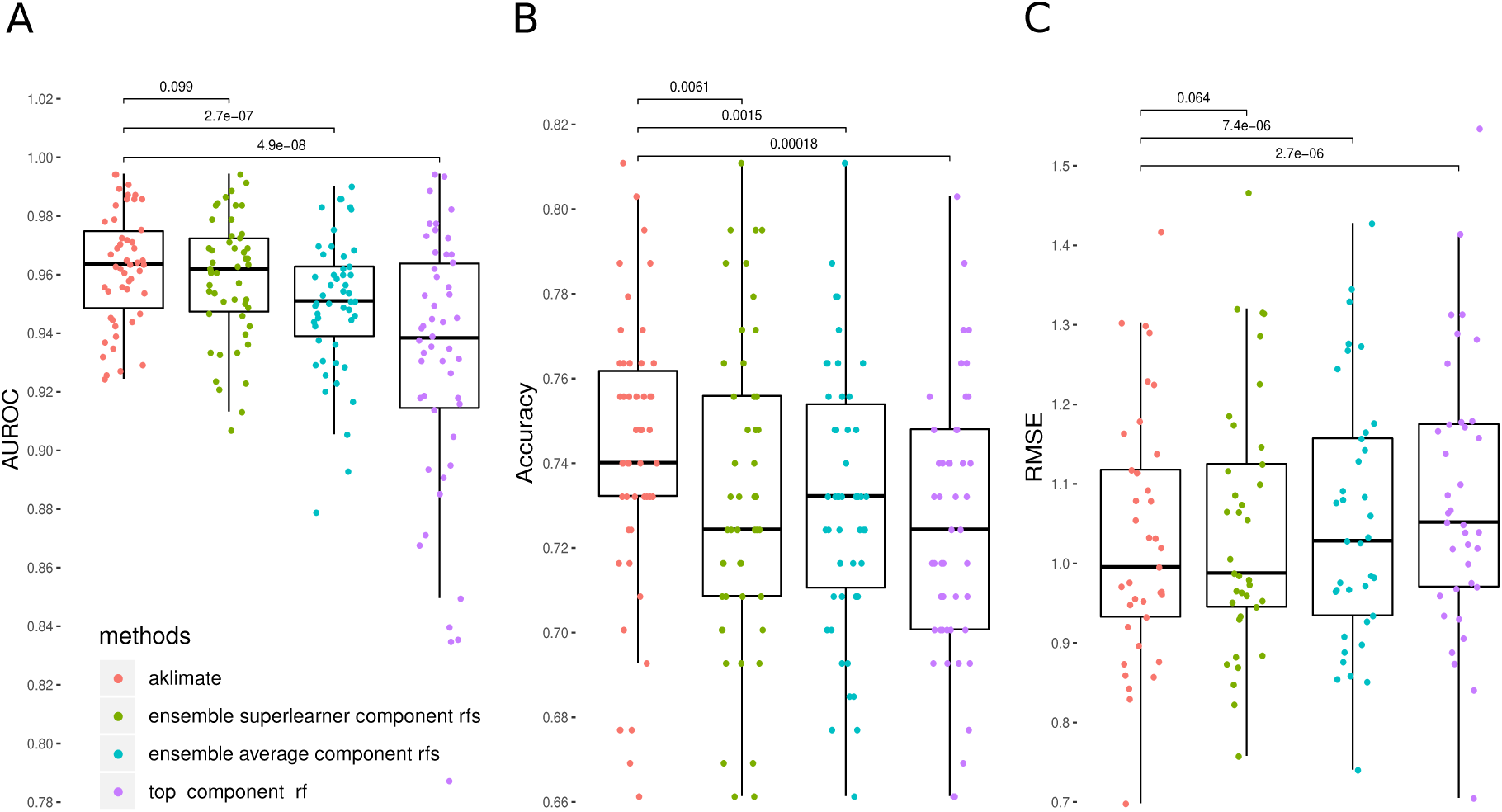
AKLIMATE versus alternative ways of ensembling/stacking component RFs. To ensure fair comparison, the component RF set for each prediction task was restricted to AKLIMATE’s best RFs **Δ**\*. Ensemble superlearner component RFs - learning a regularized linear regression on predictions from component RFs; ensemble average component RFs - taking the average of the predictions of component RFs; top component RF - only using predictions from the top ranked RF. A) UCEC TCGA MSI prediction. B) METABRIC Breast Cancer survival prediction. C) Achilles shRNA knockdown prediction. P-values for Wilcoxon signed-rank test pairwise comparisons.

**Fig. S2.**
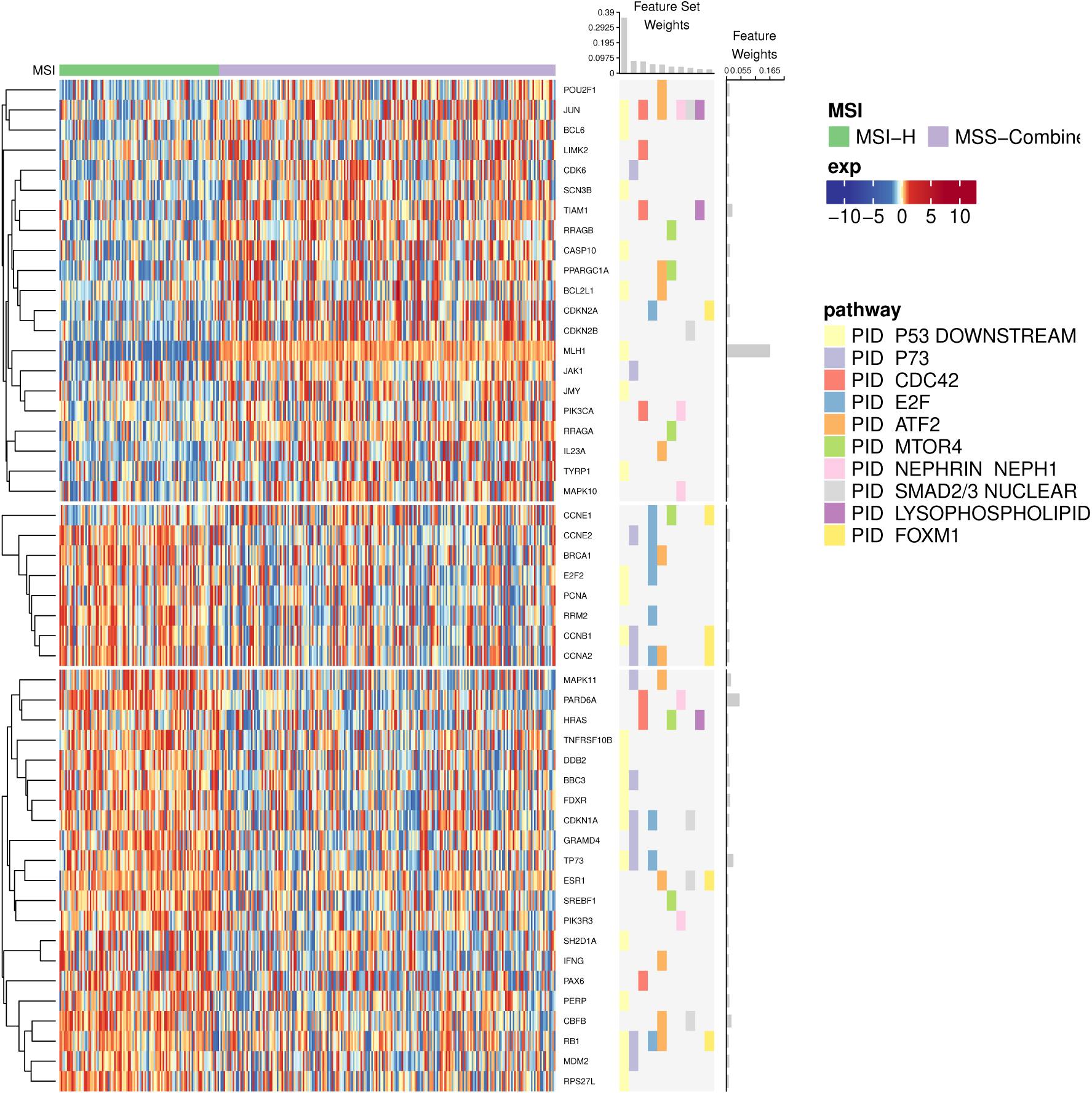
AKLIMATE results for the AKLIMATE-reduced model (using only 196 PID pathways) on UCEC TCGA cohort. Top 10 predictive feature sets and top 50 predictive features from 50 stratified 75% train/25% test splits are shown. Organized as Fig. 3.

**Fig. S3.**
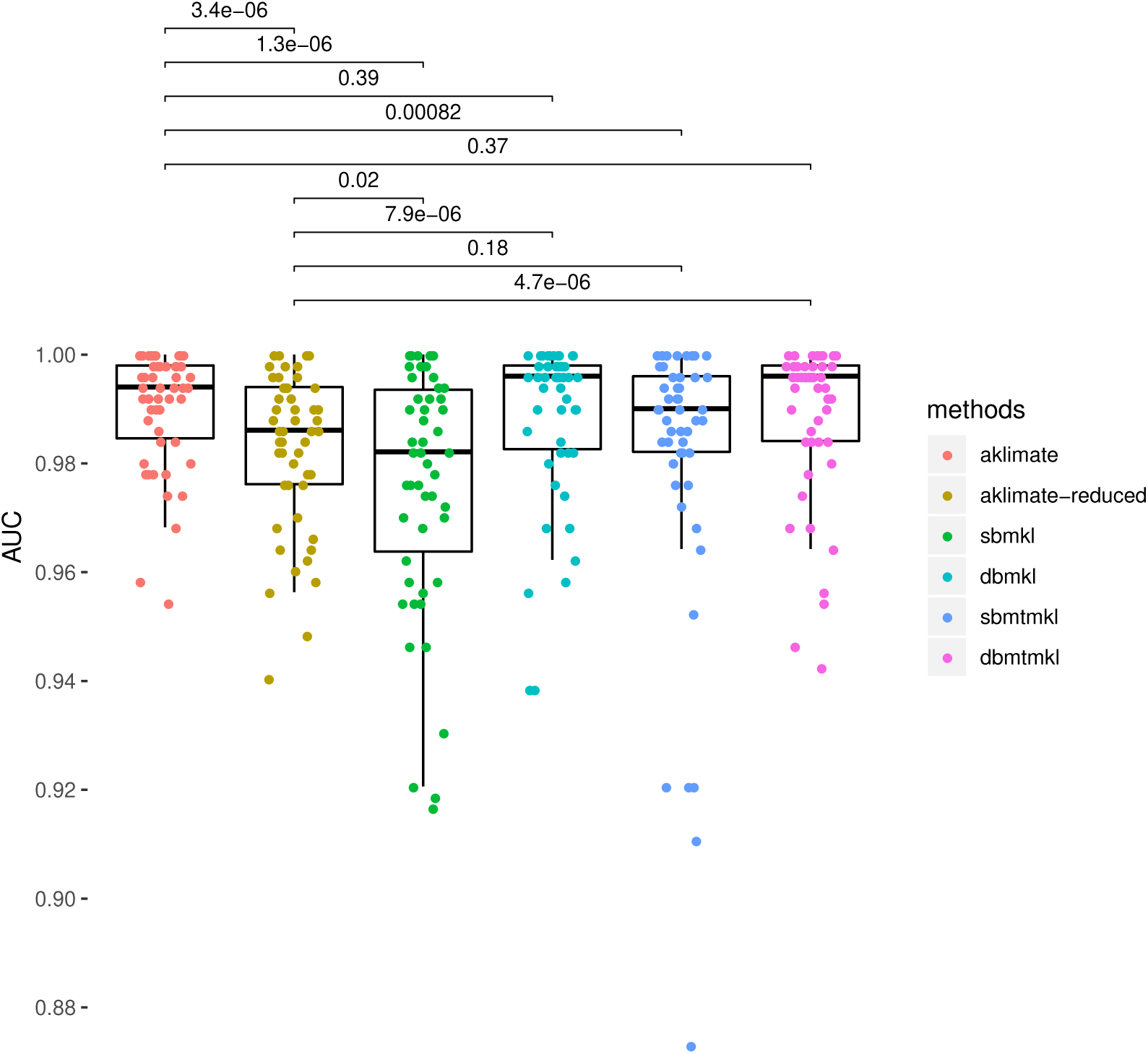
AKLIMATE performance on predicting MSI-High vs MSI-Low+MSS in TCGA COAD-READ cohort. AUC computed for 50 75%/25% stratified train/test splits. P-values for Wilcoxon signed-rank test pairwise comparisons. Methods as Fig.3.

**Fig. S4.**
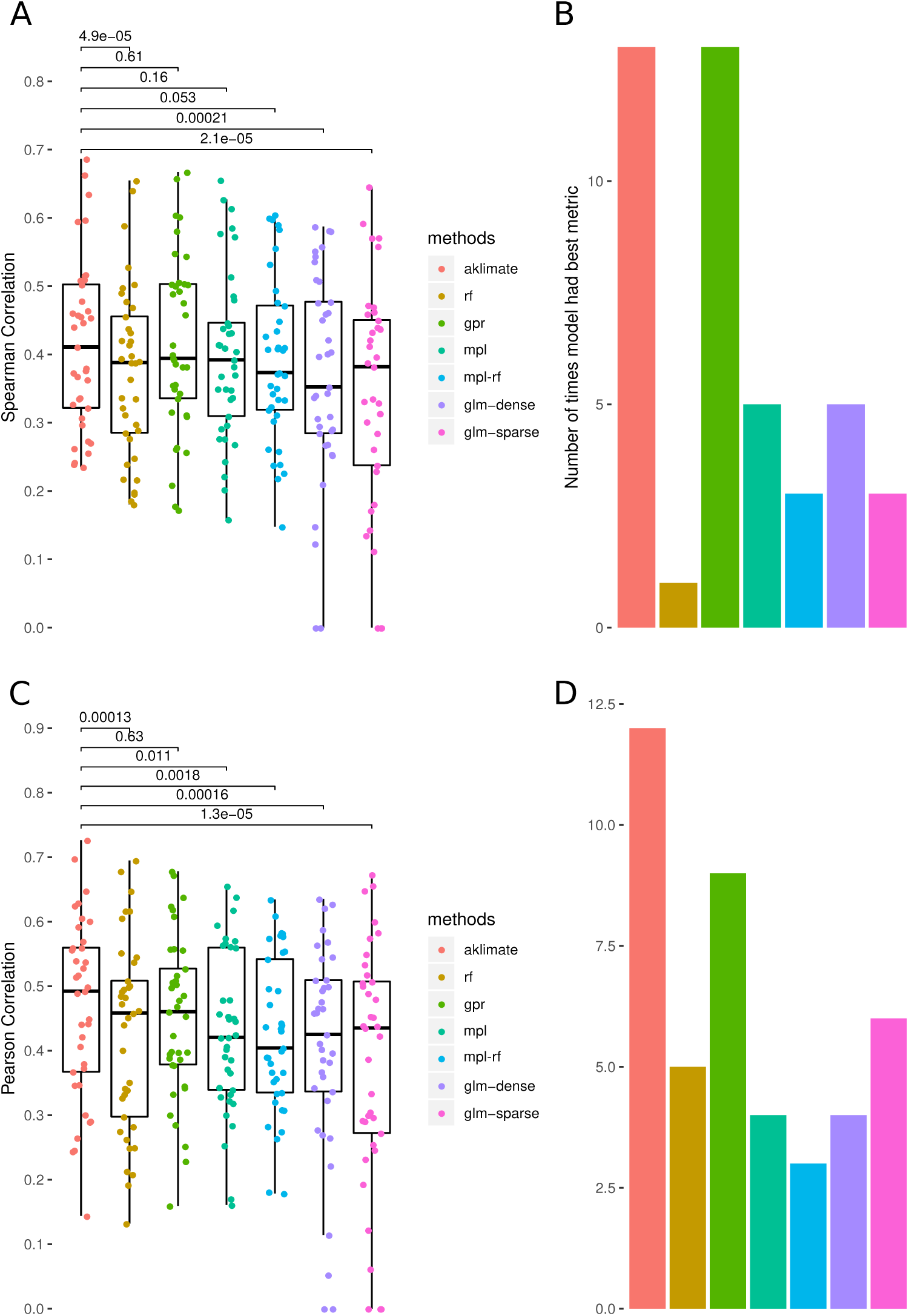
Method performance on predicting cell line viability after shRNA gene knockdowns measured by A) Spearman correlation. B) Number of times an algorithm produced the best Spearman correlation on a prediction task. C) Pearson correlation. D) Number of times an algortihm produced the best Pearson correlation on a prediction task.

**Fig. S5.**
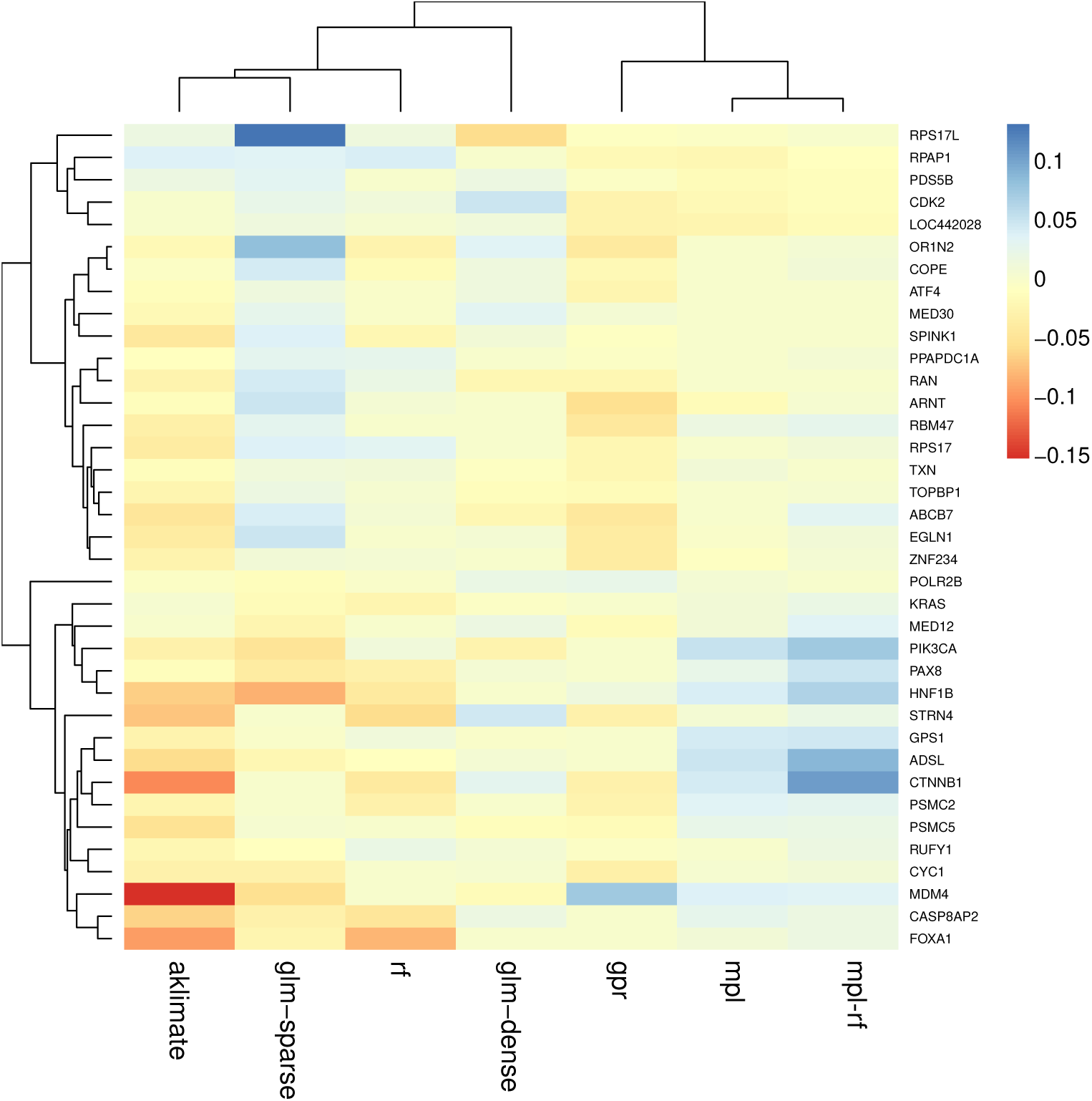
RMSE scores for 37 shRNA knockdown prediction tasks. Rows correspond to individual gene knockdowns; columns represent different methods. To highlight differential performance within each task, rows are centered by subtracting the median. Lower centered scores represent lower RMSE (better performance).

**Fig. S6.**
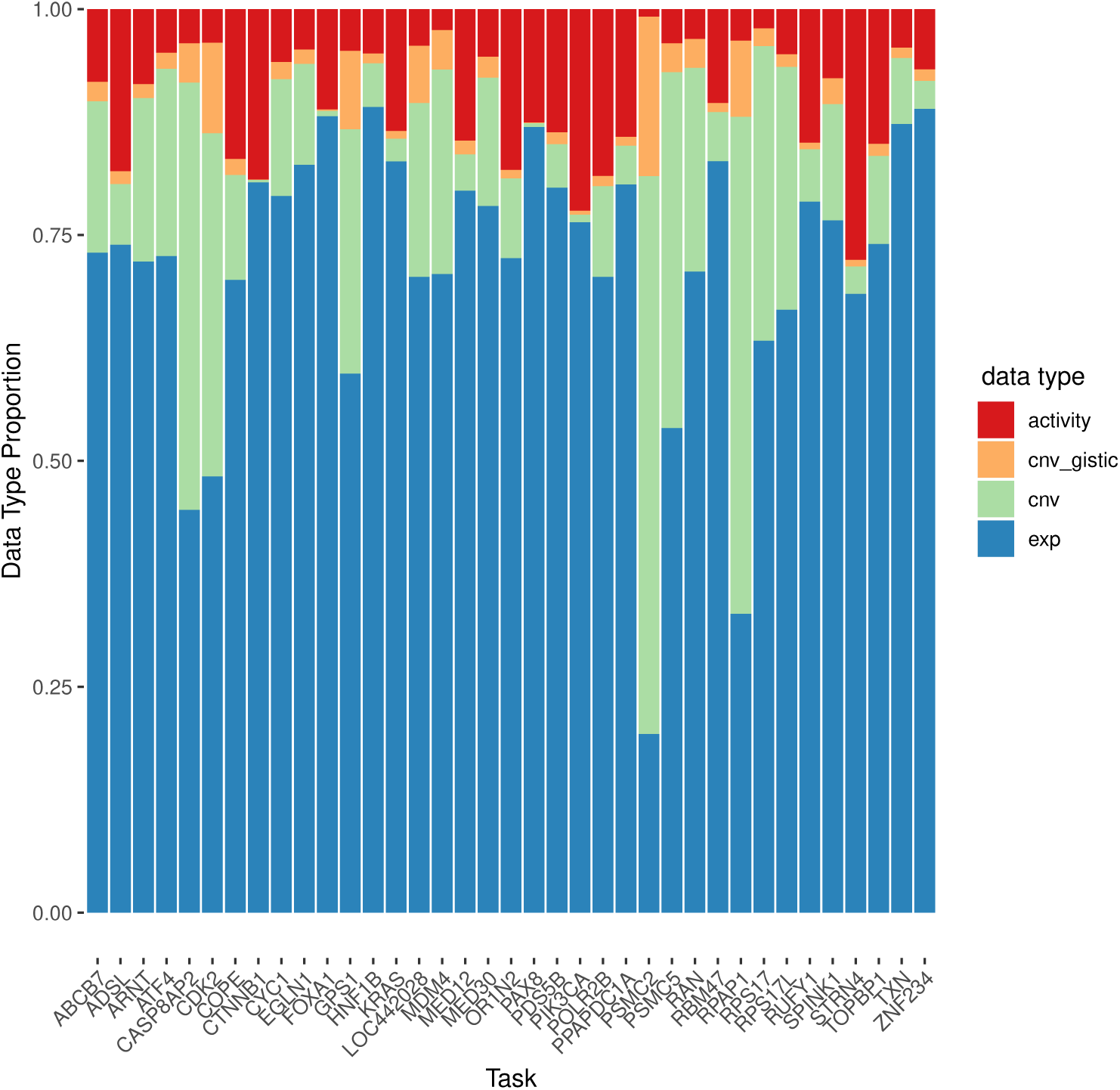
Data type importance proportions for 37 shRNA knockdown prediction tasks. Columns correspond to the relative contributions of each input data type across prediction tasks. Relative contributions were computed by summing the model-assigned feature importance scores within individual data types. Columns are normalized to sum to 1.

**Fig. S7.**
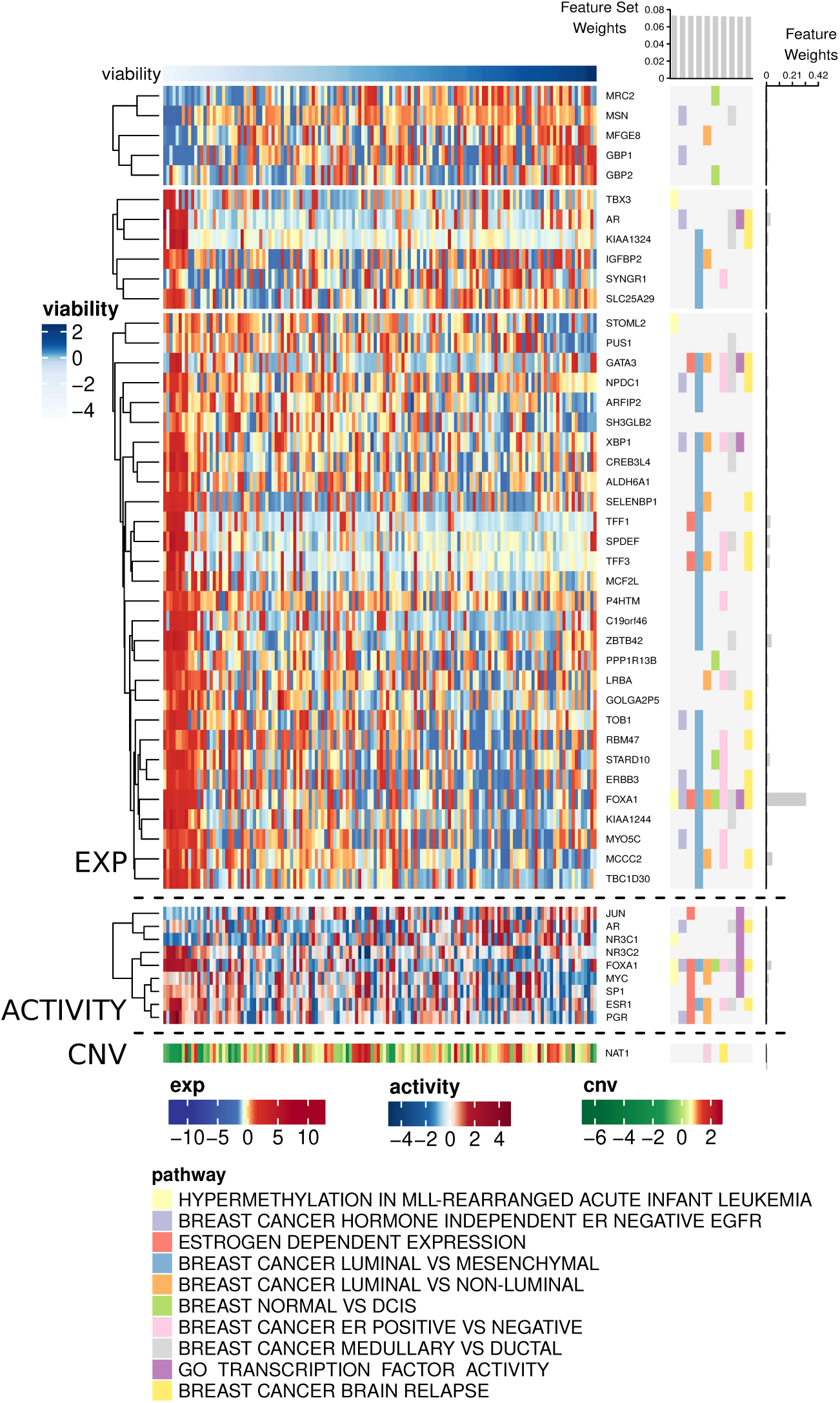
AKLIMATE results highlighting the 10 most informative feature sets and 50 most informative features for the task of predicting FOXA1 shRNA knockdown viability. Organized as Fig. 3.

**Fig. S8.**
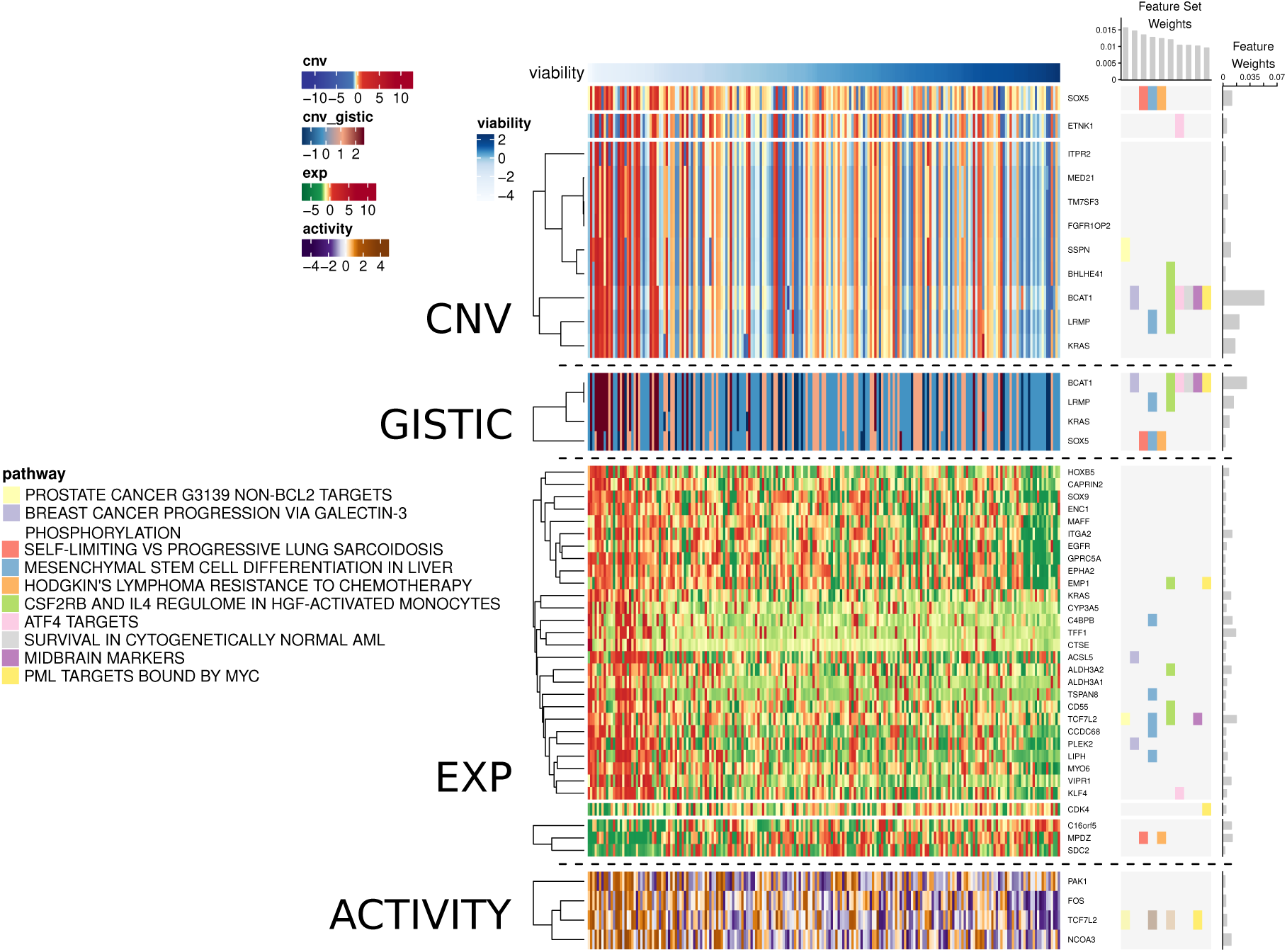
AKLIMATE results highlighting the 10 most informative feature sets and 50 most informative features for the task of predicting KRAS shRNA knockdown viability when no mutation features are used. Feature and feature set weights averaged over 10 matched stratified 80%/20% train/test splits. Organized as Fig. 3.

**Fig. S9.**
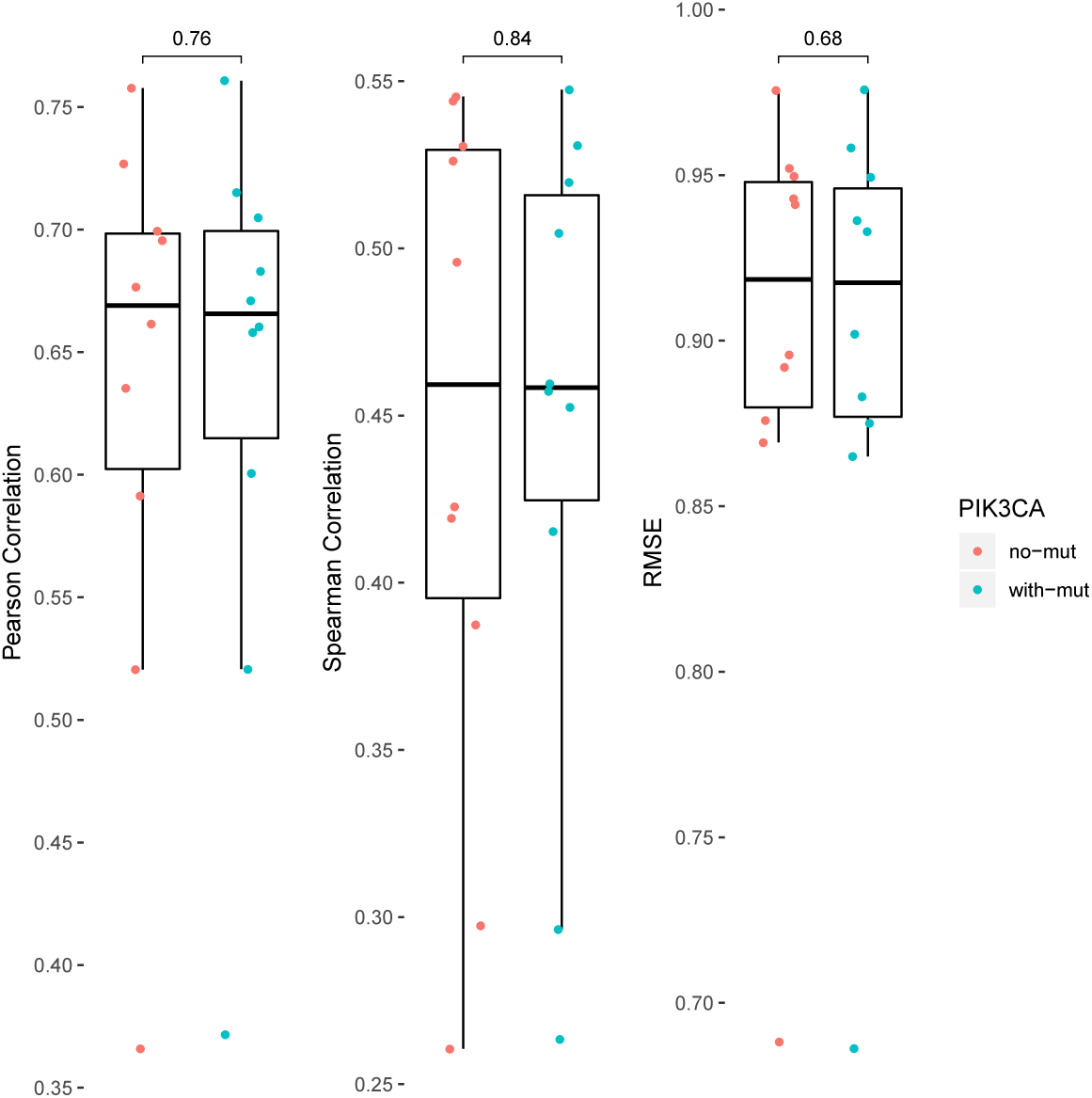
Metrics for PIK3CA AKLIMATE models with and without the use of mutational profiles for 8 key regulators. Results averaged over 10 matched stratified 80%/20% train/test splits.

**Fig. S10.**
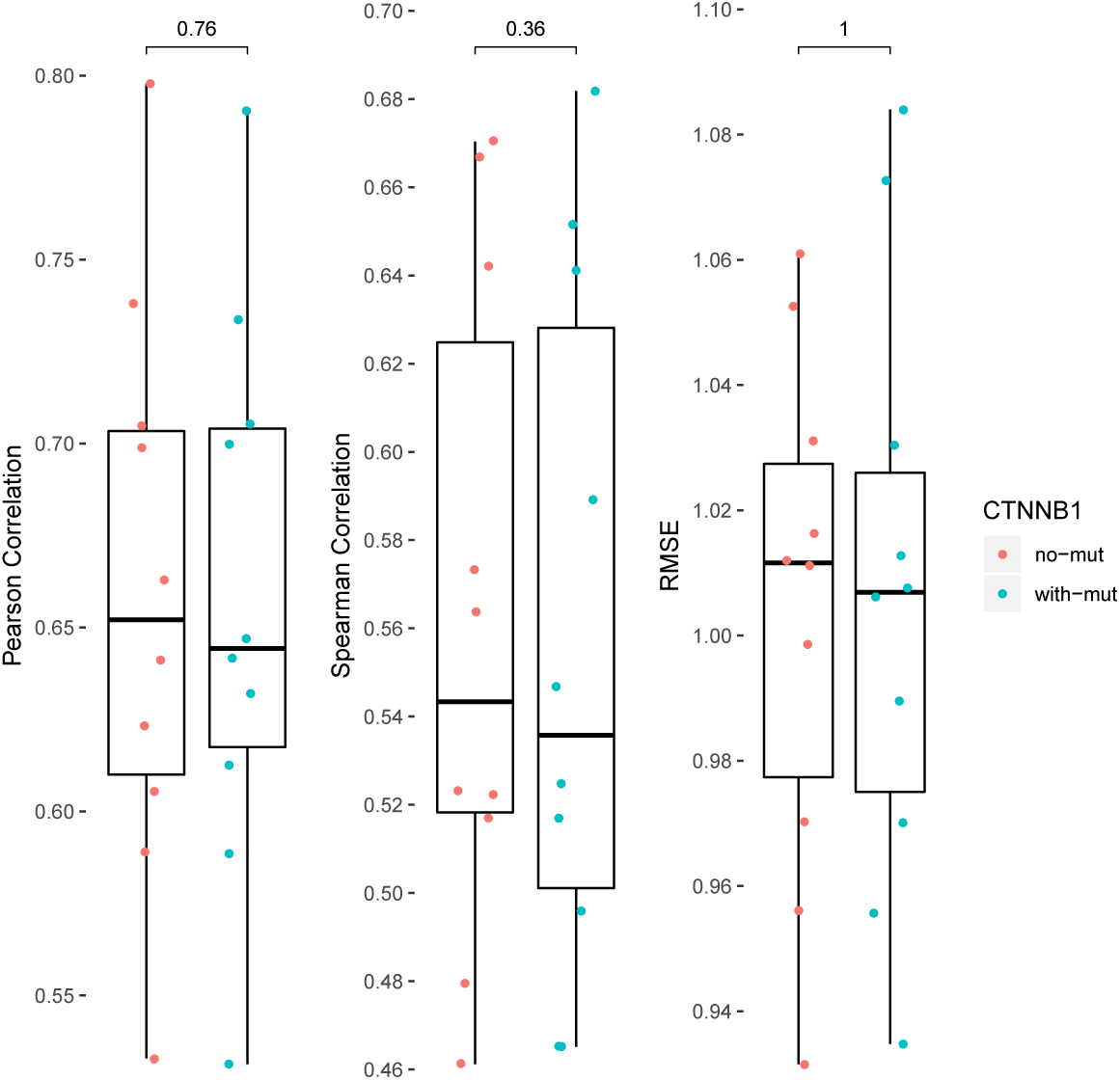
Metrics for CTNNB1 AKLIMATE models with and without the use of mutational profiles for 8 key regulators. Results averaged over 10 matched stratified 80%/20% train/test splits.

## References

[1] A. Altmann, L. Toloşi, O. Sander, and T. Lengauer. Permutation importance: a corrected feature importance measure. Bioinformatics, 26(10):1340–1347, May 2010. ISSN 1367-4803.

[2] N. Aronszajn. Theory of Reproducing Kernels. Transactions of the American Mathematical Society, 68(3):337, May 1950. ISSN 00029947.

[3] F. R. Bach, G. R. G. Lanckriet, and M. I. Jordan. Multiple kernel learning, conic duality, and the SMO algorithm. In Twenty-first international conference on Machine learning - ICML ‘04, page 6, Banff, Alberta, Canada, 2004. ACM Press.

[4] J. Barretina et al. The Cancer Cell Line Encyclopedia enables predictive modelling of anticancer drug sensitivity. Nature, 483(7391):603–607, Mar. 2012. ISSN 0028-0836.

[5] E. Bilal et al. Improving Breast Cancer Survival Analysis through Competition-Based Multidimensional Modeling. PLoS Computational Biology, 9(5):e1003047, May 2013. ISSN 1553-7358.

[6] L. Breiman. Stacked regressions. Machine Learning, 24(1):49–64, July 1996. ISSN 0885-6125, s1573-0565.

[7] L. Breiman. Some infinity theory for predictor ensembles. Technical Report 577, UC Berkeley, 2000.

[8] L. Breiman. Random Forests. Machine Learning, 45(1):5–32, Oct. 2001. ISSN 1573-0565.

[9] L. Breiman, J. H. Friedman, Olshen, R.A., and Stone, C.J. Classification and Regression Trees. Routledge, 1984. ISBN 978-1-351-46049-1.

[10] H. Cao, S. Bernard, R. Sabourin, and L. Heutte. Random forest dissimilarity based multi-view learning for Radiomics application. Pattern Recognition, 88:185–197, Apr. 2019. ISSN 0031-3203.

[11] H. Cao, S. Bernard, R. Sabourin, and L. Heutte. A Novel Random Forest Dissimilarity Measure for Multi-View Learning. arXiv:2007.02572 [cs, stat], July 2020. 2007.02572.

[12] E. G. Cerami et al. Pathway Commons, a web resource for biological pathway data. Nucleic Acids Research, 39(Database issue):D685–D690, Jan. 2011. ISSN 0305-1048.

[13] W.-Y. Cheng, T.-H. O. Yang, and D. Anastassiou. Development of a Prognostic Model for Breast Cancer Survival in an Open Challenge Environment. Science Translational Medicine, 5(181):181ra50–181ra50, Apr. 2013. ISSN 1946-6234, 1946-6242.

[14] J. C. Costello et al. A community effort to assess and improve drug sensitivity prediction algorithms. Nature Biotechnology, 32(12):1202–1212, Dec. 2014. ISSN 1546-1696.

[15] G. S. Cowley et al. Parallel genome-scale loss of function screens in 216 cancer cell lines for the identification of context-specific genetic dependencies. Scientific Data, 1:140035, Sept. 2014. ISSN 2052-4463.

[16] C. J. Creighton et al. Development of resistance to targeted therapies transforms the clinically-associated molecular profile subtype of breast tumor xenografts. Cancer research, 68(18):7493–7501, Sept. 2008. ISSN 0008-5472.

[17] C. J. Creighton et al. Molecular profiles of progesterone receptor loss in human breast tumors. Breast cancer research and treatment, 114(2):287–299, Mar. 2009. ISSN 0167-6806.

[18] A. C. Culhane et al. GeneSigDB: a manually curated database and resource for analysis of gene expression signatures. Nucleic Acids Research, 40(Database issue):D1060–D1066, Jan. 2012. ISSN 0305-1048.

[19] C. Curtis et al. The genomic and transcriptomic architecture of 2,000 breast tumours reveals novel subgroups. Nature, 486(7403):346–352, June 2012. ISSN 1476-4687.

[20] A. Davies and Z. Ghahramani. The Random Forest Kernel and other kernels for big data from random partitions. arXiv:1402.4293 [cs, stat], Feb. 2014. 1402.4293.

[21] L. Ding et al. Perspective on Oncogenic Processes at the End of the Beginning of Cancer Genomics. Cell, 173(2):305–320.e10, Apr. 2018. ISSN 0092-8674.

[22] C. Englund and A. Verikas. A novel approach to estimate proximity in a random forest: An exploratory study. Expert Systems with Applications, 39(17):13046–13050, Dec. 2012. ISSN 0957-4174.

[23] J. H. Friedman. Stochastic gradient boosting. Computational Statistics & Data Analysis, 38(4):367–378, Feb. 2002. ISSN 0167-9473.

[24] J. H. Friedman, T. Hastie, and R. Tibshirani. Regularization Paths for Generalized Linear Models via Coordinate Descent. Journal of Statistical Software, 33(1):1–22, Feb. 2010. ISSN 1548-7660.

[25] P. Geurts, D. Ernst, and L. Wehenkel. Extremely randomized trees. Machine Learning, 63(1):3–42, Apr. 2006. ISSN 0885-6125, 1573-0565.

[26] A. E. Giuliano et al. Breast Cancer—Major changes in the American Joint Committee on Cancer eighth edition cancer staging manual. CA: A Cancer Journal for Clinicians, 67(4):290–303, 2017. ISSN 1542-4863.

[27] M. Gönen. Integrating gene set analysis and nonlinear predictive modeling of disease phenotypes using a Bayesian multitask formulation. BMC Bioinformatics, 17(16):0, Dec. 2016. ISSN 1471-2105.

[28] M. Gönen and E. Alpaydın. Multiple Kernel Learning Algorithms. J. Mach. Learn. Res., 12:2211–2268, July 2011. ISSN 1532-4435.

[29] M. Gönen et al. A Community Challenge for Inferring Genetic Predictors of Gene Essentialities through Analysis of a Functional Screen of Cancer Cell Lines. Cell Systems, 5(5):485–497.e3, Nov. 2017. ISSN 2405-4712.

[30] J. L. Haybittle et al. A prognostic index in primary breast cancer. British Journal of Cancer, 45(3):361–366, Mar. 1982. ISSN 0007-0920.

[31] M. Hitchins et al. Dominantly Inherited Constitutional Epigenetic Silencing of MLH1 in a Cancer-Affected Family Is Linked to a Single Nucleotide Variant within the 5’UTR. Cancer Cell, 20(2):200–213, Aug. 2011. ISSN 1535-6108.

[32] K. A. Hoadley et al. Multiplatform Analysis of 12 Cancer Types Reveals Molecular Classification within and across Tissues of Origin. Cell, 158(4):929–944, Aug. 2014. ISSN 0092-8674, 1097-4172w.

[33] K. A. Hoadley et al. Cell-of-Origin Patterns Dominate the Molecular Classification of 10,000 Tumors from 33 Types of Cancer. Cell, 173(2):291–304.e6, Apr. 2018. ISSN 0092-8674, 1097-4172.

[34] G. A. Hobbs, C. J. Der, and K. L. Rossman. RAS isoforms and mutations in cancer at a glance. Journal of Cell Science, 129(7):1287–1292, Apr. 2016. ISSN 0021-9533.

[35] S. Huang, K. Chaudhary, and L. X. Garmire. More Is Better: Recent Progress in Multi-Omics Data Integration Methods. Frontiers in Genetics, 8, 2017. ISSN 1664-8021.

[36] N. Hunter and R. H. Borts. Mlh1 is unique among mismatch repair proteins in its ability to promote crossing-over during meiosis. Genes & Development, 11(12):1573–1582, June 1997. ISSN 0890-9369, 1549-5477.

[37] L. Jacob, G. Obozinski, and J.-P. Vert. Group lasso with overlap and graph lasso. In Proceedings of the 26th Annual International Conference on Machine Learning - ICML ‘09, pages 1–8, Montreal, Quebec, Canada, 2009. ACM Press. ISBN 978-1-60558-516-1.

[38] T. Jewison et al. SMPDB 2.0: big improvements to the Small Molecule Pathway Database. Nucleic Acids Research, 42(Database issue):D478–484, Jan. 2014. ISSN 1362-4962.

[39] R. N. Jorissen et al. DNA copy-number alterations underlie gene expression differences between microsatellite stable and unstable colorectal cancers. Clinical Cancer Research: An Official Journal of the American Association for Cancer Research, 14(24):8061–8069, Dec. 2008. ISSN 1078-0432.

[40] M. Kanehisa et al. KEGG for integration and interpretation of large-scale molecular data sets. Nucleic Acids Research, 40(D1):D109–D114, Jan. 2012. ISSN 0305-1048.

[41] M. Kim, N. Rai, V. Zorraquino, and I. Tagkopoulos. Multi-omics integration accurately predicts cellular state in unexplored conditions for Escherichia coli. Nature Communications, 7, Oct. 2016. ISSN 2041-1723.

[42] G. Kimeldorf and G. Wahba. Some results on Tchebycheffian spline functions. Journal of Mathematical Analysis and Applications, 33(1):82–95, Jan. 1971. ISSN 0022-247X.

[43] M. Kloft, U. Rückert, and P. L. Bartlett. A Unifying View of Multiple Kernel Learning. In D. Hutchison et al., editors, Machine Learning and Knowledge Discovery in Databases, volume 6322, pages 66–81. Springer Berlin Heidelberg, Berlin, Heidelberg, 2010. ISBN 978-3-642-15882-7 978-3-642-15883-4.

[44] G. R. G. Lanckriet et al. A statistical framework for genomic data fusion. Bioinformatics, 20(16):2626–2635, Nov. 2004. ISSN 1367-4803, 1460-2059.

[45] E. LeDell. Scalable Ensemble Learning and Computationally Efficient Variance Estimation. PhD thesis, University of California, Berkeley, 2015.

[46] C. Li and H. Li. Network-constrained regularization and variable selection for analysis of genomic data. Bioinformatics, 24(9):1175–1182, May 2008. ISSN 1367-4803, 1460-2059.

[47] A. Liberzon et al. Molecular signatures database (MSigDB) 3.0. Bioinformatics, 27(12): 1739–1740, June 2011. ISSN 1367-4803.

[48] G. Louppe. Understanding Random Forests: From Theory to Practice. arXiv:1407.7502 [stat], July 2014. 1407.7502.

[49] S. Mallik and Z. Zhao. Graph- and rule-based learning algorithms: a comprehensive review of their applications for cancer type classification and prognosis using genomic data. Briefings in Bioinformatics, 2019.

[50] M. Manica, J. Cadow, R. Mathis, and M. Rodríguez Martínez. PIMKL: Pathway-Induced Multiple Kernel Learning. npj Systems Biology and Applications, 5(1):1–8, Mar. 2019. ISSN 2056-7189.

[51] D. Marbach et al. Wisdom of crowds for robust gene network inference. Nat Meth, 9(8): 796–804, Aug. 2012. ISSN 1548-7091.

[52] A. A. Margolin et al. Systematic Analysis of Challenge-Driven Improvements in Molecular Prognostic Models for Breast Cancer. Science translational medicine, 5(181):181re1, Apr. 2013. ISSN 1946-6234.

[53] C. H. Mermel et al. GISTIC2.0 facilitates sensitive and confident localization of the targets of focal somatic copy-number alteration in human cancers. Genome Biology, 12 (4):R41, 2011. ISSN 1465-6906.

[54] Q. Mo et al. Pattern discovery and cancer gene identification in integrated cancer genomic data. Proceedings of the National Academy of Sciences, 110(11):4245–4250, Mar. 2013. ISSN 0027-8424, 1091-6490.

[55] S. Nembrini, I. R. König, M. N. Wright, and A. Valencia. The revival of the Gini importance? Bioinformatics, 2018.

[56] A. Y. Ng. Preventing “Overfitting” of Cross-Validation Data. In Proceedings of the Fourteenth International Conference on Machine Learning, ICML ‘97, pages 245–253, San Francisco, CA, USA, 1997. Morgan Kaufmann Publishers Inc. ISBN 978-1-55860-486-5.

[57] E. C. Polley. Super Learner In Prediction. Technical Report 266, UC Berkeley, 2010.

[58] D. Pratt et al. NDEx, the Network Data Exchange. Cell Systems, 1(4):302–305, Oct. 2015. ISSN 2405-4712.

[59] A. Rakotomamonjy, F. R. Bach, S. Canu, and Y. Grandvalet. SimpleMKL. Journal of Machine Learning Research, 9(Nov):2491–2521, 2008. ISSN ISSN 1533-7928.

[60] N. Rappoport and R. Shamir. Multi-omic and multi-view clustering algorithms: review and cancer benchmark. Nucleic Acids Research, 46(20):10546–10562, Nov. 2018. ISSN 1362-4962.

[61] A. G. Robertson et al. Integrative Analysis Identifies Four Molecular and Clinical Subsets in Uveal Melanoma. Cancer Cell, 32(2):204–220.e15, Aug. 2017. ISSN 1535-6108, 1878-3686.

[62] M. Sandri and P. Zuccolotto. A Bias Correction Algorithm for the Gini Variable Importance Measure in Classification Trees. Journal of Computational and Graphical Statistics, 17(3):611–628, Sept. 2008. ISSN 1061-8600, 1537-2715.

[63] C. F. Schaefer et al. PID: the Pathway Interaction Database. Nucleic Acids Research, 37 (Database issue):D674–679, Jan. 2009. ISSN 1362-4962.

[64] B. Scholkopf and A. J. Smola. Learning with Kernels: Support Vector Machines, Regularization, Optimization, and Beyond. MIT Press, Cambridge, MA, USA, 2001. ISBN 978-0-262-19475-4.

[65] E. Scornet. Random Forests and Kernel Methods. IEEE Transactions on Information Theory, 62(3):1485–1500, Mar. 2016. ISSN 0018-9448.

[66] J. A. Seoane, I. N. M. Day, T. R. Gaunt, and C. Campbell. A pathway-based data integration framework for prediction of disease progression. Bioinformatics, 30(6):838–845, Mar. 2014. ISSN 1367-4803, 1460-2059.

[67] D. D. Shao et al. ATARiS: Computational quantification of gene suppression phenotypes from multisample RNAi screens. Genome Research, 23(4):665–678, Apr. 2013. ISSN 1088-9051.

[68] J. Shawe-Taylor and N. Cristianini. Kernel Methods for Pattern Analysis. Cambridge University Press, 2004.

[69] A. Sokolov et al. Pathway-Based Genomics Prediction using Generalized Elastic Net. PLOS Computational Biology, 12(3):e1004790, Mar. 2016. ISSN 1553-7358.

[70] G. Stolovitzky, D. Monroe, and A. Califano. Dialogue on Reverse-Engineering Assessment and Methods. Annals of the New York Academy of Sciences, 1115(1):1–22, 2007. ISSN 1749-6632.

[71] C. Strobl, A.-L. Boulesteix, A. Zeileis, and T. Hothorn. Bias in random forest variable importance measures: Illustrations, sources and a solution. BMC Bioinformatics, 8(1): 25, Jan. 2007. ISSN 1471-2105.

[72] A. Subramanian et al. Gene set enrichment analysis: A knowledge-based approach for interpreting genome-wide expression profiles. Proceedings of the National Academy of Sciences of the United States of America, 102(43):15545–15550, Oct. 2005.

[73] T. Suzuki and M. Sugiyama. Fast learning rate of multiple kernel learning: Trade-off between sparsity and smoothness. The Annals of Statistics, 41(3):1381–1405, June 2013. ISSN 0090-5364.

[74] T. Suzuki and R. Tomioka. SpicyMKL: a fast algorithm for Multiple Kernel Learning with thousands of kernels. Machine Learning, 85(1-2):77–108, Oct. 2011. ISSN 0885-6125, 1573-0565.

[75] The HPN-DREAM Consortium et al. Inferring causal molecular networks: empirical assessment through a community-based effort. Nature Methods, 13(4):310–318, Apr. 2016. ISSN 1548-7091, 1548-7105.

[76] R. Tibshirani. Regression Shrinkage and Selection via the Lasso. Journal of the Royal Statistical Society. Series B (Methodological), 58(1):267–288, 1996.

[77] R. Tomioka and T. Suzuki. Sparsity-accuracy trade-off in MKL. arXiv:1001.2615 [stat], Jan. 2010. 1001.2615.

[78] V. J. Uzunangelov. Prediction of cancer phenotypes through the integration of multi-omic data and prior information. PhD thesis, UC Santa Cruz, 2019.

[79] A. W. v. d. Vaart, S. Dudoit, and M. J. v. d. Laan. Oracle inequalities for multi-fold cross validation. Statistics & Decisions, 24(3):351–371, 2006.

[80] M. van der Laan and S. Dudoit. Unified Cross-Validation Methodology For Selection Among Estimators and a General Cross-Validated Adaptive Epsilon-Net Estimator: Finite Sample Oracle Inequalities and Examples. Technical Report 130, University of California, Berkeley U.C. Berkeley Division of Biostatistics Working Paper Series, 2003.

[81] M. J. van der Laan, E. C. Polley, and A. E. Hubbard. Super Learner. Statistical Applications in Genetics and Molecular Biology, 6(1), Jan. 2007. ISSN 1544-6115, 2194-6302.

[82] Q. Wan and R. Pal. An Ensemble Based Top Performing Approach for NCI-DREAM Drug Sensitivity Prediction Challenge. PLoS ONE, 9(6), June 2014. ISSN 1932-6203.

[83] J. N. Weinstein et al. The cancer genome atlas pan-cancer analysis project. Nature genetics, 45(10):1113, 2013.

[84] D. H. Wolpert. Stacked Generalization. Neural Networks, 5:241–259, 1992.

[85] M. N. Wright and A. Ziegler. ranger: A Fast Implementation of Random Forests for High Dimensional Data in C++ and R. Journal of Statistical Software, 77(1):1–17, Mar. 2017. ISSN 1548-7660.

[86] M. Yuan and Y. Lin. Model selection and estimation in regression with grouped variables. Journal of the Royal Statistical Society: Series B (Statistical Methodology), 68(1):49–67, Feb. 2006. ISSN 1369-7412, 1467-9868.

[87] H. Zou and T. Hastie. Regularization and variable selection via the elastic net. Journal of the Royal Statistical Society: Series B (Statistical Methodology), 67(2):301–320, Apr. 2005. ISSN 1467-9868.

[88] K. Zuberi et al. GeneMANIA Prediction Server 2013 Update. Nucleic Acids Research, 41(W1):W115–W122, July 2013. ISSN 0305-1048.

